# Identifying strategies to target the metabolic flexibility of tumours

**DOI:** 10.1101/2020.01.06.896571

**Authors:** Andrés Méndez-Lucas, Wei Lin, Paul C. Driscoll, Nathalie Legrave, Laura Novellasdemunt Vilaseca, Chencheng Xie, Mark Charles, Zena Wilson, Neil P. Jones, Stephen Rayport, Manuel Rodríguez-Justo, Vivian Li, James I. MacRae, Nissim Hay, Xin Chen, Mariia Yuneva

## Abstract

Plasticity of cancer metabolism can be a major obstacle for efficient targeting of tumour-specific metabolic vulnerabilities. Here, we identify and quantify the compensatory mechanisms following the inhibition of major pathways of central carbon metabolism in c-MYC-induced liver tumours. We find that glutaminase isoform Gls2, expressed in normal liver, compensates for the deletion of Gls1 isoform expressed in tumours. Inhibiting both glutaminases significantly delays tumourigenesis but does not completely block glutamine catabolism through the Krebs cycle. We reveal that glutamine catabolism is then driven by amidotransferases. Consistently, the synergistic effect of glutaminase and amidotransferase inhibitors on proliferation of mouse and human tumour cells is observed *in vitro* and *in vivo*. Furthermore, when Gls1 is deleted the Krebs cycle activity and tumour formation can also be significantly affected if glycolysis is co-inhibited (Gls1^KO^/*Hk2^KO^*). Finally, the inhibition of either serine (*Psat1*^KO^) or fatty acid (*Fasn*^KO^) biosynthesis can be compensated by uptake of circulating nutrients. Thus, removing these nutrients from the diet produces synergistic effects on suppression of tumourigenesis. These results highlight the high flexibility of tumour metabolism and demonstrate how targeting compensatory mechanisms can improve a therapeutic outcome.

## Introduction

Metabolism of tumour cells is different from metabolism of cells in a parental tissue. This change is thought to favour a maximized use of the resources with an adequate energy balance that allows cancer cells to survive and proliferate in a competitive tumour microenvironment^1, 2^. This metabolic rewiring of tumours can impose metabolic dependencies which can be exploited to develop therapeutic approaches ^3^. However, to date only few existing strategies targeting tumour-specific metabolic vulnerabilities have proven useful ^3^. One of the underlying reasons may be metabolic plasticity of cancer cells that allows for compensatory adaptations. Another factor could be the prediction of metabolism-based therapeutic targets based on experiments utilizing *in vitro* culture conditions, which significantly affects metabolism of initial tumour-derived cells^4^, and do not recapitulate tumour heterogeneity and complex inter-tumour and tumour-host interactions^5^.

One of the major regulators of cellular metabolism during proliferation and development is proto-oncogene c-MYC (hereforth termed MYC), which is also one of the most frequently dysregulated lesions in human cancers^6, 7^. The difficulty of targeting MYC itself^8^ highlights the need to uncover and exploit therapeutic targets amongst its downstream effectors, including metabolic pathways. Using a genetically engineered mouse model of liver cancer^9^ we and others have demonstrated that MYC-induced liver tumours have increased catabolism of both glucose and glutamine in comparison with the normal liver^10, 11^. These metabolic changes are associated with a repression of some liver-specific enzymes as well as the expression of enzyme isoforms that are not expressed in the normal liver^10–12^, including glutaminase 1 (GLS1) that catalyses the first step of glutamine catabolism, and the HK2 isoform of the first enzyme of glycolysis, hexokinase. Since both glucose and glutamine fuel pathways that are vital for proliferation and survival of cells, including the Krebs cycle, or serine, nucleotide, and lipid biosynthesis^4, 13–16^, several enzymes controlling these pathways have been considered as attractive therapeutic targets^4, 10, 17–21^. However, the inability to reach a complete inhibition of tumourigenesis in any of these experiments suggests that tumours engage mechanisms of compensation that allow them to survive and proliferate despite decreased activity of these enzymes. Although the understanding of the complexity of factors determining tumour metabolic vulnerabilities and flexibilities is emerging^1, 22–25^, the mechanisms of metabolic resistance of tumours *in vivo* still remain largely unknown. Here we have combined metabolomics, genomics and pharmacological approaches to demonstrate the high flexibility of tumour metabolism, and how blocking the mechanisms of compensation can lead to stronger inhibition of tumour growth *in vivo*.

## Results

### Major pathways of glucose and glutamine catabolism are increased in MYC-induced liver tumours

We have previously demonstrated that glucose and glutamine catabolism are increased in the MYC-induced mouse liver tumour model, LAP-tTA/TRE-MYC^10^. To further evaluate which pathways fuelled by glucose and glutamine are affected in these tumours, we performed *in vivo* metabolic studies using either boluses or infusions of stable isotope-labelled nutrients, [U-^13^C]-glucose and [U-^13^C]-glutamine, in tumour-bearing mice^9^. Mice kept on a doxycycline diet to suppress MYC expression were used as controls. A short time bolus allowed us to evaluate the capacity of tissues to take up and catabolize these nutrients. Intravenous infusions (3 hours) allowed us to measure a tracer incorporation into various metabolites, including lipids, and to calculate the steady state partial contribution of glucose and glutamine to different pathways. The tissue metabolites were extracted and analysed by gas chromatography mass-spectrometry (GC-MS) and ^13^C-^1^H-2D-HSQC nuclear magnetic resonance (NMR). A bolus of [U-^13^C]-glucose resulted in increased enrichment of lactate and citrate (Fig. 1a) while a bolus of [U-^13^C]-glutamine resulted in increased enrichment of citrate, malate, fumarate and oxaloacetate-derived aspartate in tumours in comparison with the normal livers (Fig. 1b). These results suggested that glucose catabolism through glycolysis and catabolism of both glucose and glutamine through the Krebs cycle are increased on MYC-induced liver tumours in comparison with normal tissue counterparts consistent with our previously published results^10, 12^.

**Fig. 1.**
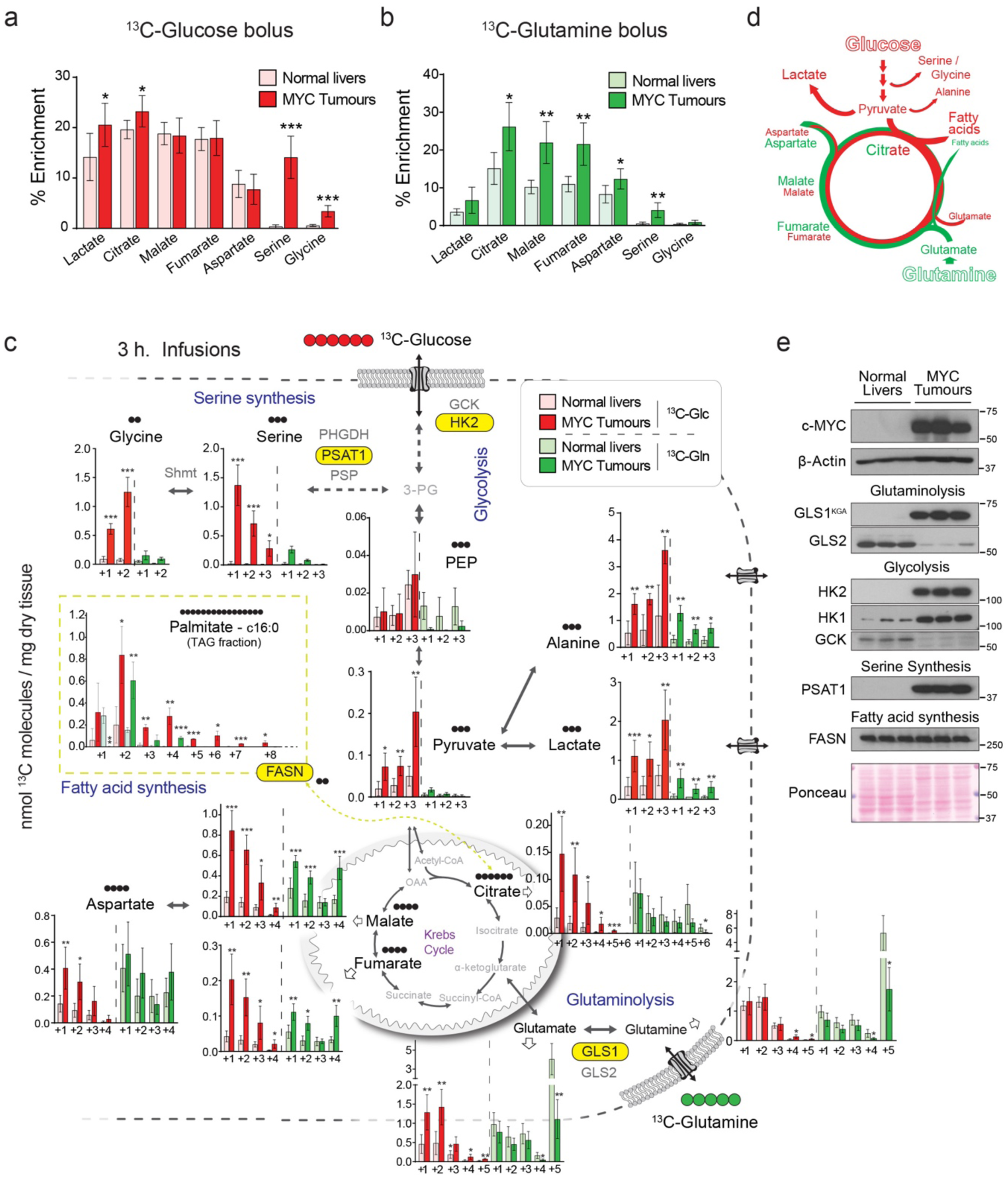
*In vivo* metabolic tracing using ^13^C-labeled glucose and glutamine reveals upregulated pathways in MYC-driven liver tumours. (a and b) ^13^C-enrichment of the indicated metabolites extracted from either normal livers or MYC-driven liver tumours after either a bolus of [U-^13^C]-glucose (a, n=5-6) or [U-^13^C]-glutamine (b, n=4-5) measured by GC-MS. Tumours were initiated in LAP-tTA/TRE-MYC mice by weaning them into regular chow. Mice kept on a doxycycline-containing diet were used as controls. (c) Quantification of the [U-^13^C]-glucose and [U-^13^C]-glutamine carbon incorporation into some of the key intermediates of the central carbon metabolism in MYC-driven liver tumours and control normal livers after 3 hours of infusion. Data are presented as the mean ± S.D., n = 5. *, p < 0.05; **, p < 0.01; ***, p < 0.001 relative to the same isotopologue in normal livers infused with the same label. The content of ^13^C-labeled palmitic acid from a triglyceride pool is compared for tumours and corresponding adjacent livers. Glutamine labelling is estimated from quantification of its spontaneous product pyroglutamate, and glutamate from the non-pyroglutamate fraction. (d) Diagram depicting the relative contribution of glucose (in red) and glutamine (in green) to the different pathways studied (e), Western blot comparing MYC-driven tumours and normal livers for the expression levels of key enzymes involved in the metabolic pathways depicted in E. See also Extended data Fig 1 and 2.

This conclusion was further confirmed by the results of 3-hour infusions of either [U-^13^C]-glucose or [U-^13^C]-glutamine. The partial contribution of glucose and glutamine to the pools of most of the Krebs cycle intermediates at steady-state remained similar to that observed in normal livers, except for ^13^C-glucose-derived citrate, the entry point of glucose carbons into the Krebs cycle (Extended Data Fig. 1a,b,d,e,g). However, substantially increased pools of the Krebs cycle intermediates in tumours (Extended Data Fig. 1c,f) suggested that under steady-state conditions higher catabolism of glucose and glutamine was required in tumours to label them to the same extent as in normal tissues. Indeed, the pools of ^13^C-glucose-derived pyruvate, lactate, alanine, citrate and glutamate, and the ^13^C-carbon pools of malate, fumarate, and aspartate derived from either [U-^13^C]-glucose or [U-^13^C]-glutamine were significantly increased in tumours, in comparison with the normal liver (Fig. 1c). Concomitant decrease of tumoural ^13^C-glutamate and ^13^C-glutamine pools suggested their fast catabolism and increased anaplerosis. Labelling in all carbon positions of pyruvate, lactate and alanine from both ^13^C-glucose and ^13^C-glutamine in tumours also suggested malic enzyme-driven pyruvate cycling. The extent of metabolite enrichment either from glucose or glutamine demonstrated that they are major sources for the generation of the Krebs cycle intermediates in the MYC-induced liver tumours, with a substantial contribution of these nutrients to the Krebs cycle in both tumours and livers (30-40% of total isotope enrichment). These results demonstrate a simultaneous enhancement of glucose and glutamine catabolism through the Krebs cycle in tumours but which still preserves the partial contribution of both nutrients.

Together with fuelling the Krebs cycle, the increase in glucose and glutamine catabolism in MYC-induced liver tumours could be required to support the activity of other pathways essential for tumourigenesis, such as amino acid and lipid biosynthesis (Fig. 1d). Indeed, we observed a marked increase in serine and glycine production from [U-^13^C]-glucose (Fig. 1a and Extended Data Fig. 1b) and alpha-^15^N-glutamine (Extended Data Fig. 1h) in tumours, and an accumulation of both metabolites (Extended Data Fig. 1c,f), compared to normal livers. Interestingly, as we have described before, the main serine isotopologues observed during U-^13^C]-glucose infusion were +1 and +2 and not +3, which can be produced due to carbon exchange during glycine synthesis and one carbon metabolism^26^. We also observed a strong increase in the incorporation of both glucose and glutamine carbons into tumour fatty acids (Fig. 1c and Extended Data Fig. 1i) consistent with the observed accumulation of neutral lipids in tumours (Extended Data Fig. 1j).

The increased activities of glycolysis, the Krebs cycle, and serine and lipid biosynthesis in tumours were associated with an elevated expression of enzymes or a change in enzyme isoform patterns (Fig. 1e and Extended Data Fig. 2a,b) consistent with previous results published by us and others^10, 11^. These changes included increased expression of *Hk2* and decreased expression of glucokinase (*Gck*, or Hexokinase 4; glycolysis) and increased expression of *Gls1* and decreased expression of the *Gls2* isoform of glutaminase (glutaminolysis). In addition, the expression of all enzymes of the serine biosynthesis pathway was higher in tumours compared with normal livers, most notably phosphoserine aminotransferase (*Psat1*) (Fig. 1e and Extended Data Fig. 2b). Finally, although the mRNA expression levels of enzymes involved in some normal liver functions, such as gluconeogenesis, were suppressed in the tumours, the expression of some lipogenic enzymes, such as fatty acid synthase (FASN), remained high (Fig. 1e and Extended Data Fig. 2b).

To evaluate whether the increased activity of these major pathways of glucose and glutamine catabolism is required for MYC-induced liver tumourigenesis, we used the Albumin-CreER^T2^ mouse line to perform hepatocyte-specific knock-out of *Hk2*, *Gls1*, *Psat1*, and *Fasn* genes at the time of tumour initiation (Extended Data Fig. 2c). To induce MYC-driven tumours in this case we used hydrodinamics-based transfection of a MYC-encoding plasmid^12, 27^. Since ectopic expression of Cre recombinase has been previously shown to exacerbate MYC-induced apoptosis and reduce the efficiency of MYC-induced tumourigenesis^28^, we also combined the ectopic expression of MYC with the expression of MCL1, another gene frequently overexpressed in human liver tumours (Extended Data Fig. 2d). We verified the hepatocyte-specific origin of MYC-induced tumours by adeno-associated virus-mediated lineage tracing (Extended Data Fig. 2e-h)^29^.

### Glutaminases are not the only drivers of glutamine catabolism

Liver-specific knock-out of *Gls1* (*Gls1*^fl/fl^+Alb-CreER^T2^+Tamoxifen, *Gls1*^KO^; Figure 2a) significantly increased the latency of MYC-induced liver tumours (Figure 2b), confirming the key role of *Gls1* for MYC-induced tumourigenesis^18^. *Gls1*^KO^ tumours from animals injected with [U-^13^C]-glutamine had significantly higher levels of glutamine (Figure 2c), and a lower incorporation of glutamine-derived carbons into the Krebs cycle intermediates and amino acids (Figure 2d), than control tumours (Alb-CreER^T2^+Tamoxifen; CT). However, the total levels of the Krebs cycle intermediates were not affected upon *Gls1* KO (Extended Data Figure 3a). These data demonstrated that GLS1 plays a significant role in glutaminolysis in MYC-induced liver tumours, but also showed that in the absence of GLS1 tumours were still able to develop, and catabolize a substantial amount of glutamine. We next investigated which alternative enzymes could be responsible for this capability.

**Fig. 2.**
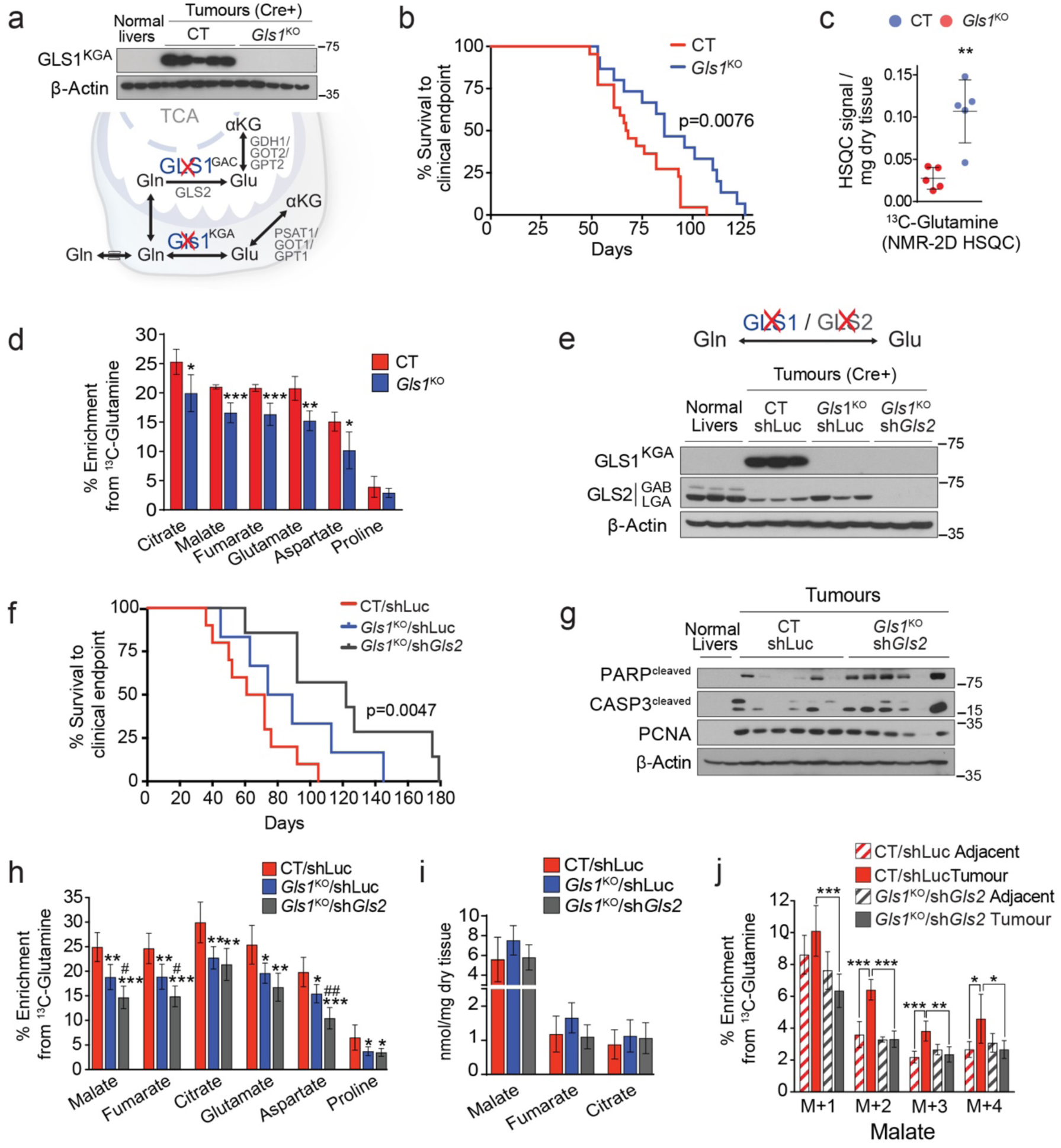
Ablation of glutaminases impairs MYC-induced tumourigenesis and reveals a contribution of a glutaminase-independent glutamine catabolism. (a-d) Inhibiting *Gls1* expression in MYC-driven tumours decreases glutamine catabolism and tumour burden: Tumours were induced by hydrodynamics-based transfection of MYC/MCL1 2 weeks after tamoxifen-induced liver-specific CRE activation in *Gls1*^fl/fl^/Alb-CreER^T2^/Rosa26eYFP and Alb-CreER^T2^/Rosa26eYFP mice. (a) Top, western blot demonstrating the absence of GLS1 in *Gls1*^KO^ tumours. Bottom, glutaminases catalyse the conversion of glutamine to glutamate, a key point of amino acid metabolism; (b) Kaplan-Meier Survival Curve (CT n=21; *Gls1*^KO^ n=15); (c) Glutamine levels in CT and *Gls1*^KO^ tumours (n=5 per group, NMR); (d) ^13^C-enrichment from [U-^13^C]-glutamine in the indicated metabolites of CT and *Gls1*^KO^ tumours after [U-^13^C]-glutamine boluses (n= 5 per group, GC-MS). (e-j) *Gls2* knock-down in *Gls1*^KO^ tumours increases the repressive effect on tumour glutaminolysis and tumour burden: (e) Western blot demonstrating shRNA-mediated reduction of GLS2 protein levels in tumours; (f) Kaplan-Meier Survival Curve (CT/shLuc n=10, *Gls1*^KO^/shLuc n=6, *Gls1*^KO^/sh*Gls2* n=7); (g) Western blot showing the protein levels of cleaved PARP, cleaved Caspase 3 and PCNA in the indicated tissues. β-Actin was used as a loading control; (h) ^13^C-enrichment of the indicated metabolites after a [U-^13^C]-glutamine bolus (n= 6, 5, 6, group labels as above); (i) Total levels of the Krebs cycle intermediates (n=6, 5, 6, group labels as above; GC-MS); (j) ^13^C-enrichment of malate isotopologues in the experiment shown in Fig. 2h. See also Extended data Fig 2, 3 and 4.

GLS2, whose expression was decreased in MYC-induced tumours in comparison with the normal liver, appeared to be expressed in *Gls1*^KO^ tumours (Figure 2e and Extended Data Figure 3b). *Gls2* knock-down in *Gls1* wild type tumours, using short hairpin RNA (shRNA) interference (Extended Data Figure 3c), had no impact either on tumourigenesis or on incorporation of [U-^13^C]-glutamine-derived carbons into downstream intermediates (Extended Data Figure 3d,e). This demonstrated that GLS1 is the major glutaminase isoform supporting glutamine catabolism in these tumours. To test whether GLS2 supports glutamine catabolism in the absence of GLS1, we inhibited *Gls2* expression in *Gls1*^KO^ tumours (*Gls1*^KO^/sh*Gls2*) (Figure 2e). *Gls1^KO^/shGls2* tumours were compared to *CT/shLuc* and *Gls1KO/shLuc* tumours expressing shRNA against luciferase. *Gls1^KO^/shGls2* combination resulted in a more significant delay in tumourigenesis (Figure 2f), which was associated with increased levels of apoptosis markers cleaved caspase 3 and poly-ADP ribose polymerase (PARP) and reduced levels of a proliferation marker, proliferating cell nuclear antigen (PCNA), when compared to *CT/shLuc* (Figure 2g). These data suggested that the growth of *Gls1*^KO^/sh*Gls2* tumours was inhibited by an increased level of tumour cell apoptosis and reduced proliferation rate, and not only at the tumour initiation stage. *Gls1*^KO^/sh*Gls2* tumours showed a trend towards even higher accumulation of glutamine in comparison with the one observed in *Gls1^KO^* tumours (Extended Data Figure 3f) and a further suppression of the incorporation of [U-^13^C]-glutamine-derived carbons into the Krebs cycle and amino acids compared to *Gls1* deletion alone (Figure 2h and Extended Data Figure 3g,h). Nevertheless, the levels of the Krebs cycle intermediates were still not significantly affected by inhibiting the expression of both glutaminases (Figure 2i).

Interestingly, while the levels of the Krebs cycle intermediates were preserved in both *Gls1*^KO^ and *Gls1*^KO^/sh*Gls2* tumours, the total levels of non-essential amino acids (NEAA’s), whose synthesis depends on glutamine, including glutamate, alanine and aspartate, showed a tendency to decrease in both types of tumours in comparison with their control counterparts (Extended Data Figure 4a,b). To explore a possible mechanism for the delayed formation of *Gls1*^KO^ and *Gls1*^KO^/sh*Gls2* tumours, we compared metabolic profiles of cells derived from *CT/shLuc*, *Gls1KO/shLuc* and *Gls1^KO^/shGls2* tumours (MYC-driven hepatocellular carcinoma (HCC) cells; HCC^MYC^). Consistent with the *in vivo* phenotype, HCC^MYC^ - *Gls1*^KO^*/shLuc* cells proliferated slower than HCC^MYC^ cells derived from *CT/shLuc* tumours and this effect was significantly exacerbated in *Gls1*^KO^/sh*Gls2* tumour cells (Extended Data Fig. 4c). Importantly, the levels of the Krebs cycle intermediates, malate and fumarate, were only decreased in *Gls1*^KO^/sh*Gls2* cells and the levels of both aspartate and alanine were decreased in *Gls1KO/shLuc* and *Gls1*^KO^/sh*Gls2* cells (Extended Data Fig. 4d). The extent of these changes correlated with the extent of proliferation delay observed in these cell lines. Interestingly the levels of citrate were not affected (Extended Data Fig. 4d) possibly being supported by glucose catabolism. These results suggested that decreasing glutamine catabolism into the Krebs cycle may affect its biosynthetic capacity supporting tumour cell proliferation. Consistently with this hypothesis, the presence of all NEAA’s in the media improved significantly slowed proliferation of HCC^MYC^-*Gls1*^KO^/sh*Gls2* cells (Extended Data Fig. 4c). The fact that the extent of the decrease in the levels of the Krebs cycle intermediates and amino acids was much more significant *in vitro* than *in vivo* could be due to the uptake of the amino acids by tumours from the bloodstream. To test if this is indeed the case, we administered mice bearing either CT/shLuc or *Gls1*^KO^/sh*Gls2* tumours with a bolus of ^15^N-alanine, the second most abundant amino acid in serum, after glutamine. Interestingly, both types of tumours were not only able to uptake alanine, but also converted it into glutamate, and consequently the rest of NEAA’s by using the amino group of alanine, with an initial step catalyzed by alanine aminotransferases (Extended Data Fig. 4f). Together these data suggest that limiting a biosynthetic capacity of the Krebs cycle may be one of the factors determining the effect of glutaminase inhibition on tumour progression. Furthermore, the uptake and catabolism of circulating amino acids can support amino acid pools in tumours and may constitute one of the factors of resistance against glutaminase inhibition.

Although *Gls1*^KO^/sh*Gls2* tumours had significantly decreased flux of glutamine into the Krebs cycle, they were still able to incorporate approximately half of the glutamine-derived carbons into the Krebs cycle intermediates, compared to control tumours, and still at the levels comparable to the adjacent livers (Fig. 2h and Extended Data Fig. 3h). The direct glutamine catabolism in *Gls1*^KO^/sh*Gls2* tumours was reflected by the presence of a substantial fraction of the +4 isotopologue of malate derived from [U-^13^C]-glutamine (Fig. 2j). We therefore hypothesised that enzymes other than glutaminases could be responsible for the production of glutamate from glutamine and subsequent flux into the Krebs cycle, and could be potentially co-targeted together with glutaminase inhibition for a therapeutic intervention. Indeed, several other enzymes can utilize glutamine as an amide donor and generate glutamate ^30^ (Fig. 3a). These include enzymes involved in asparagine, nucleotide, glucosamine, NADH, and Aminoacyl-tRNA biosynthesis (Fig. 3b). Interestingly, we found that the expression of most of these enzymes was upregulated in liver tumours driven by MYC (Fig. 3c). Furthermore, the analysis of publicly-available data demonstrated that the expression of GLS1 correlates with the expression of amidotransferases in human HCC (Extended Data Fig. 5a) indicating that GLS1 inhibitors may not be efficient as a single therapy in human patients and should be combined with inhibitors of amidotransferases.

**Fig. 3.**
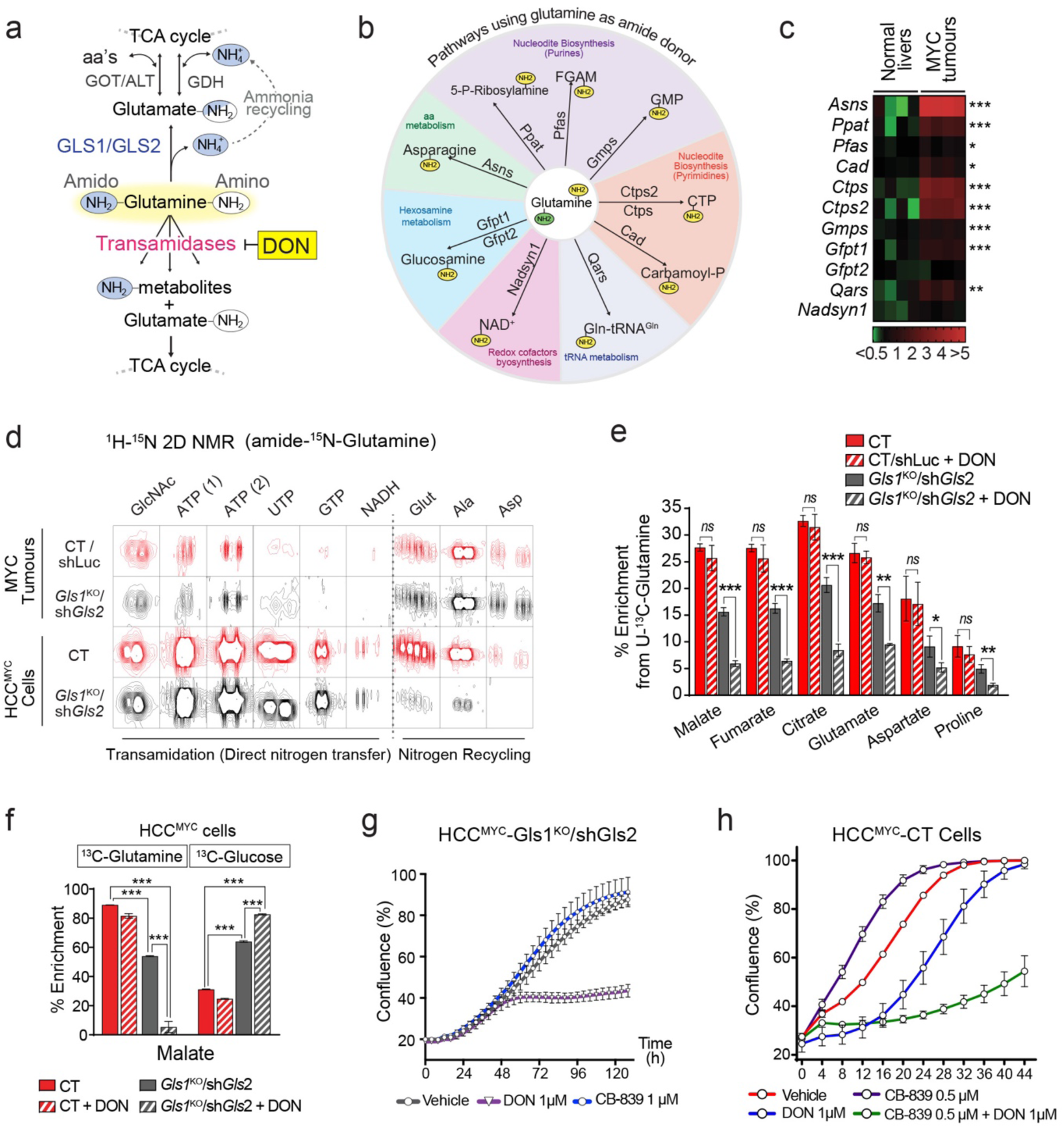
Enzymes that use glutamine as an amide donor sustain glutamine catabolism when glutaminase activity is inhibited. (a) The fate of the amide group of glutamine. (b) A diagram depicting the enzymes that can utilize glutamine as an amide donor, representing the pathways involved in the glutaminase-independent glutamine catabolism. (c) Heat map of transcriptome data showing the gene expression of enzymes that use glutamine as an amide donor (n=4) in tumours initiated in LAP-tTA/TRE-MYC mice by weaning them into regular chow. Mice kept on a doxycycline-containing diet are used as controls. (d) Representative images of ^1^H-^15^N 2D-HMBC NMR spectra of tumours from mice infused with amido-^15^N-glutamine, or cells derived from CT and *Gls1*^KO^/sh*Gls2* tumours incubated with amido-^15^N-glutamine for 48 hours, showing nitrogen incorporation into the indicated metabolites. (e) Effect of the glutamine antagonist and transamidase inhibitor DON (50 mg/kg, 4 hours) on the ^13^C-incorporation into metabolites in CT/shLuc and *Gls1*^KO^/sh*Gls2* tumours from mice administered with a [U-^13^C]-glutamine bolus (n=3 per group; GC-MS). (f) The effect of DON on the incorporation of either [U-^13^C]-glutamine or [U-^13^C]-glucose-derived carbons into malate in CT and *Gls1*^KO^/sh*Gls2* tumour cells (3 hr, n=3, GC-MS). (g and h) Combination of glutaminase inhibition and DON on proliferation of HCC^MYC^ cells: either HCC^MYC^-*Gls1*^KO^/sh*Gls2* (g) or HCC^MYC^-CT cells (h) were treated with the indicated concentrations of CB-839 and DON, or their indicated combinations. Growth was monitored in an IncuCyte Live-Cell analysis system. Representative curves from three independent experiments are shown. Data are presented as the mean ± S.D., *, *p* < 0.05; **, *p*< 0.01; ***, *p* < 0.001; two-tailed t-test. See also Figures S2, S5 and S6.

To investigate which of these pathways beyond glutaminase activity can contribute to the synthesis of glutamate from glutamine, we infused mice bearing either control or *Gls1*^KO^/sh*Gls2* tumours with amido-^15^N-glutamine. We also used amido-^15^N-glutamine-containing media to incubate the cells isolated from these tumours (HCC^MYC^-CT and HCC^MYC^-*Gls1*^KO^/sh*Gls2*). ^15^N-^1^H-2D-HMBC nuclear magnetic resonance (NMR) and liquid-chromatography mass-spectrometry (LC-MS) analysis of labelled tumours and cells revealed that glutamine-derived ^15^N was incorporated into nucleotides (both purines, AXP and GXP, and pyrimidines, UXP, TXP, and CXP), NADH, and glucosamine, among other metabolites (Fig. 3d and Extended Data Fig. 5b,c). Although during a direct synthesis of all non-essential amino acids from glutamine their nitrogen is derived from the glutamine’s amino group, we observed the incorporation of the amido-^15^N-glutamine-derived nitrogen into the amino acids (Fig. 3d and Extended Data Fig. 5b,c). This outcome suggested ammonia recycling, probably through the described glutamate dehydrogenase (GDH) reverse reaction^31^. This labelling pattern was dramatically reduced in *Gls1*^KO^/sh*Gls2* cells, consistent with the lack of ammonia generation, but relatively preserved in *Gls1*^KO^/sh*Gls2* tumours probably due to the intake of the labelled amino acids that were detected in serum (Extended Data Fig. 5d). To demonstrate that the sum of these transamidase reactions could be feeding the Krebs cycle with glutamine-derived carbons in the absence of glutaminase activity, we treated tumour-bearing mice with 6-diazo-5-oxo-L-norleucine (DON), a glutamine antagonist, and pan-amidotransferase inhibitor that blocks the incorporation of ^15^N-amido group of glutamine into different metabolites, including nucleotides (Extended Data Fig. 5e).

Consistent with our hypothesis, a single dose of DON prior to the administration of [U-^13^C]-glutamine substantially suppressed the incorporation of the label into the Krebs cycle intermediates and amino acids in *Gls1*^KO^/sh*Gls2* tumours (Fig. 3e), increased the levels of ^13^C-glutamine while decreasing the levels of ^13^C-glutamate (Extended Data Fig. 6a) and reduced the levels of alanine, malate, and fumarate (Extended Data Fig. 6b,c). Interestingly, DON minimally affected glutamine-derived anaplerosis in control tumours (Fig. 3e), suggesting that glutaminases are relatively insensitive to DON, and that only the combination of DON treatment with the inhibition of glutaminase activity can completely suppress glutamine catabolism. Consistently with the *in vivo* results, DON almost completely ablated glutamine utilization in HCC^MYC^-*Gls1*^KO^/sh*Gls2* cells, but had a much lesser impact on control cells (Fig. 3f and Extended Data Fig. 6d,e). Additionally, DON treatment further increased the contribution of glucose to the Krebs cycle in HCC^MYC^ -*Gls1*^KO^/sh*Gls2* cells making it virtually the sole carbon source (Fig. 3f and Extended Data Fig. 6e). These results demonstrate that in the absence of glutaminase activity the action of amidotransferases is required to sustain the glutamine catabolism into the Krebs cycle to support its activity as well as amino acid biosynthesis. Consistent with this, treatment with DON inhibited the proliferation of HCC^MYC^ cells only in combination with either the inhibition of glutaminase isoform expression (*Gls1*^KO^/sh*Gls2*, Fig. 3g) or GLS1 inhibitor, CB-839 (Fig. 3h and Extended Data Fig. 6f). The proliferation of HCC^MYC^-*Gls1*^KO^/sh*Gls2* cells treated with DON and HCC^MYC^ CT cells treated with a combination of CB-839 and DON was rescued by either the addition of a mix of NEAA’s or a combination of four amino acids absent in DMEM, i.e. alanine, aspartate, proline and asparagine (AAAP, Extended Data Fig. 6g,h). The combination of DON and CB-839 also showed a synergistic effect in the human cancer cell line HepG2 (Fig. 6i). These data demonstrate that decreasing the Krebs cycle anaplerosis and amino acid biosynthesis plays a substantial role in suppressing tumour cell proliferation downstream of combined inhibition of glutaminases and amidotransferases. Altogether, our data demonstrate that high levels of glutamine catabolism are required for MYC-induced tumourigenesis *in vivo*, and that deleting *Gls1* can be partially compensated for by the presence of *Gls2*. Our data also demonstrate that a significant proportion of glutamine-derived glutamate can be produced by enzymes that utilize glutamine as the amide donor, fuelling the Krebs cycle.

Next, we evaluated if the synergistic interaction between glutaminases and amidotransferases in supporting the activity of the Krebs cycle and tumourigenesis also exists in other tumour types. We tested this hypothesis in *in vitro* and *in vivo* models of colorectal cancer, a cancer type in which GLS1 plays a role in supporting tumour growth^32^, with MYC being one of the regulators of the colon cancer metabolic reprograming^33^. The combination of CB-839/DON combination had a significantly stronger inhibitory effect on proliferation of human colon cell line, HCT116 (Fig. 4a), and proliferation and colony formation of primary mouse Apc^min/+^ tumour organoids (Fig. 4b and c), than CB-839 or DON alone. Importantly, this effect was also observed in primary tumour organoids derived from two human patients with colorectal cancer (Fig. 4e-f), a model demonstrating a strong predictive power for the efficiency of therapeutic options in humans^34^. Finally, to evaluate if combining Gls1 and amidotransferase inhibitors is feasible and efficient *in vivo*, we treated SCID mice bearing HCT116 xenografts with a combination of an orally bioavailable selective Gls1 inhibitor (Compound 27)^35^ and DON. Consistent with the *in vitro* results, the strongest suppression of tumour growth was achieved by combining the two inhibitors (Figure 4g). Accordingly, the combination also had the strongest effect on the levels of the Krebs cycle intermediates and their labelling from ^13^C-glutamine (Fig. 4b and c). Interestingly, citrate levels were maintained, which reinforced the role of glucose as the main contributor to the citrate pool. Importantly, these results also demonstrated that DON alone does not produce a suppression of glutamine carbon entry into the Krebs cycle. DON treatment resulted in increased levels of aspartate, which may be a result of asparagine synthetase inhibition and may increase glutamine availability for glutaminases. Together these data confirm that inhibiting both glutaminases and other enzymes that use glutamine as an amide donor is required for the maximal inhibition of glutamine catabolism in tumours.

**Fig. 4.**
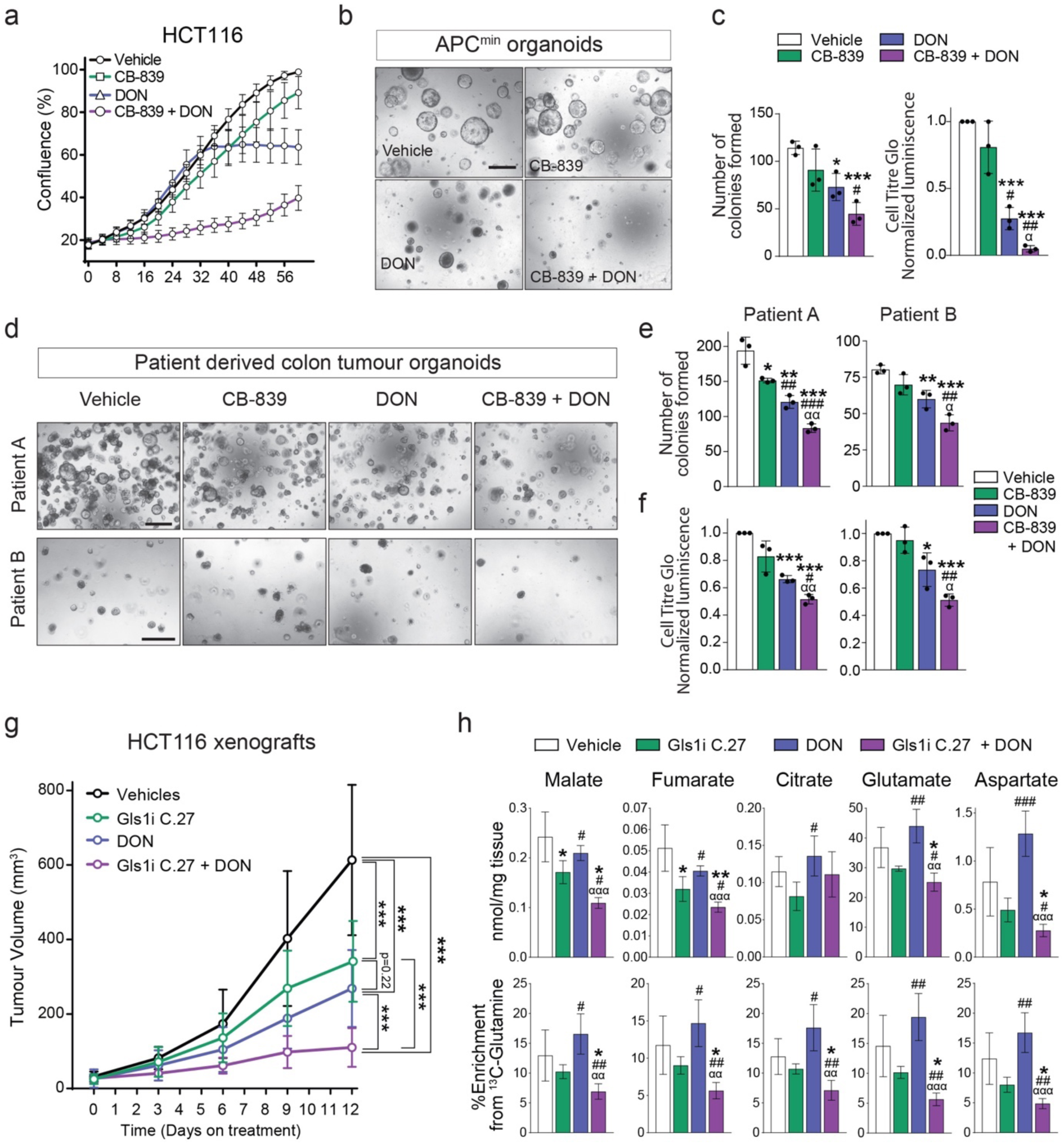
Inhibiting glutaminase and amidotransferases has a synergistic effect on mouse and human cell proliferation *in vitro* and *in vivo*. (a) HCT116 cells treated with DON and/or CB-839 at the indicated concentrations. Representative curves from three independent experiments are shown. (b) Representative photos of colony formation assay of organoids derived from APC^min^ tumours after 7 days in culture with the indicated treatments. Scale bar, 500 µm. (c) Left panel: Quantitation of the organoid formation assay in Fig. 4b. Data are mean ± S.D from three independent experiments. Right panel: Cell Titer Glo luciferase assay of APC^min^ organoids treated with the indicated treatments. Data represents three independent experiments with three replicates each. (d) Representative images of colony formation assay of two patient derived organoids after 19 days in culture (Patient A) and 10 days in culture (Patient B) with the indicated treatments. Scale bar, 500 µm. (e) Quantitation of the organoid formation assay in Fig. 4d. (f) Cell Titer Glo luciferase assay of patient derived organoids in Fig. 4d treated with the indicated treatments. In e and f, data represents three independent experiments with three replicates each. (g) The effect of combining a glutaminase inhibitor and DON on the progression of HCT116 xenografts. Mice were injected with 5 million cells and treated with 100 mg/Kg of the glutaminase inhibitor Compound 27, p.o. daily, and/or 25 mg/kg of DON every 72 hours i.p. (Vehicle n=13; Compound 27 n=10; DON n=11; Compound 27+DON n=10). (h) Total levels (top) and ^13^C-enrichment of the indicated metabolites after a [U-^13^C]-glutamine bolus in the HCT116 xenograft tumours from mice of the experiment shown in g (n= 4, GC-MS); Data are presented as the mean ± S.D., *, *p* < 0.05; **, *p*< 0.01; ***, *p* < 0.001; two-tailed t-test (c, e, f,,h), and One-Way ANOVA (g). In (h) *, # and α represent statistical significance of the indicated group versus vehicle, Gls1i C.27, or DON, respectively.

### Co-targeting glycolysis and glutaminolysis efficiently inhibits tumourigenesis

Our results demonstrated that the pools of the Krebs cycle metabolites were preserved in *Gls1*^KO^/sh*Gls2* tumours (Fig. 2h), and the glucose contribution to the Krebs cycle was increased in *Gls1*^KO^/sh*Gls2* cells in comparison with CT cells, and even more so when inhibiting glutaminase expression was combined with DON treatment (Fig. 3f and Extended Data Fig. 6d,e). We next hypothesized that concomitantly inhibiting glucose and glutamine catabolism will inhibit the Krebs cycle activity and may affect MYC-induced tumourigenesis. Increased glycolysis in MYC-induced liver tumours was associated with increased expression of *Hk2*^10^ (Fig. 1e), whose deletion was previously shown to have a profound effect on mammary gland and lung tumourigenesis^17^. First, we tested whether increased glucose catabolism was required for MYC-induced liver tumourigenesis. *Hk2* deletion at the time of tumour initiation (*Hk2*^KO^; Fig. 5a) did not affect the tumour latency (Fig. 5b), probably due to the observed compensatory re-expression of glucokinase (Fig. 5a). Nevertheless, *Hk2*^KO^ tumours showed decreased catabolism of glucose into lactate and the Krebs cycle intermediates after a [U-^13^C]-glucose bolus (Fig. 5c,d and Extended Data Fig. 7a). However, only the levels of citrate, the entry point of glucose into the Krebs cycle, were decreased but not the levels of other Krebs cycle intermediates (Extended Data Fig. 7b). Altogether these data suggested that glutaminolysis could be compensating for a decreased glycolytic flux and vice versa.

**Fig. 5.**
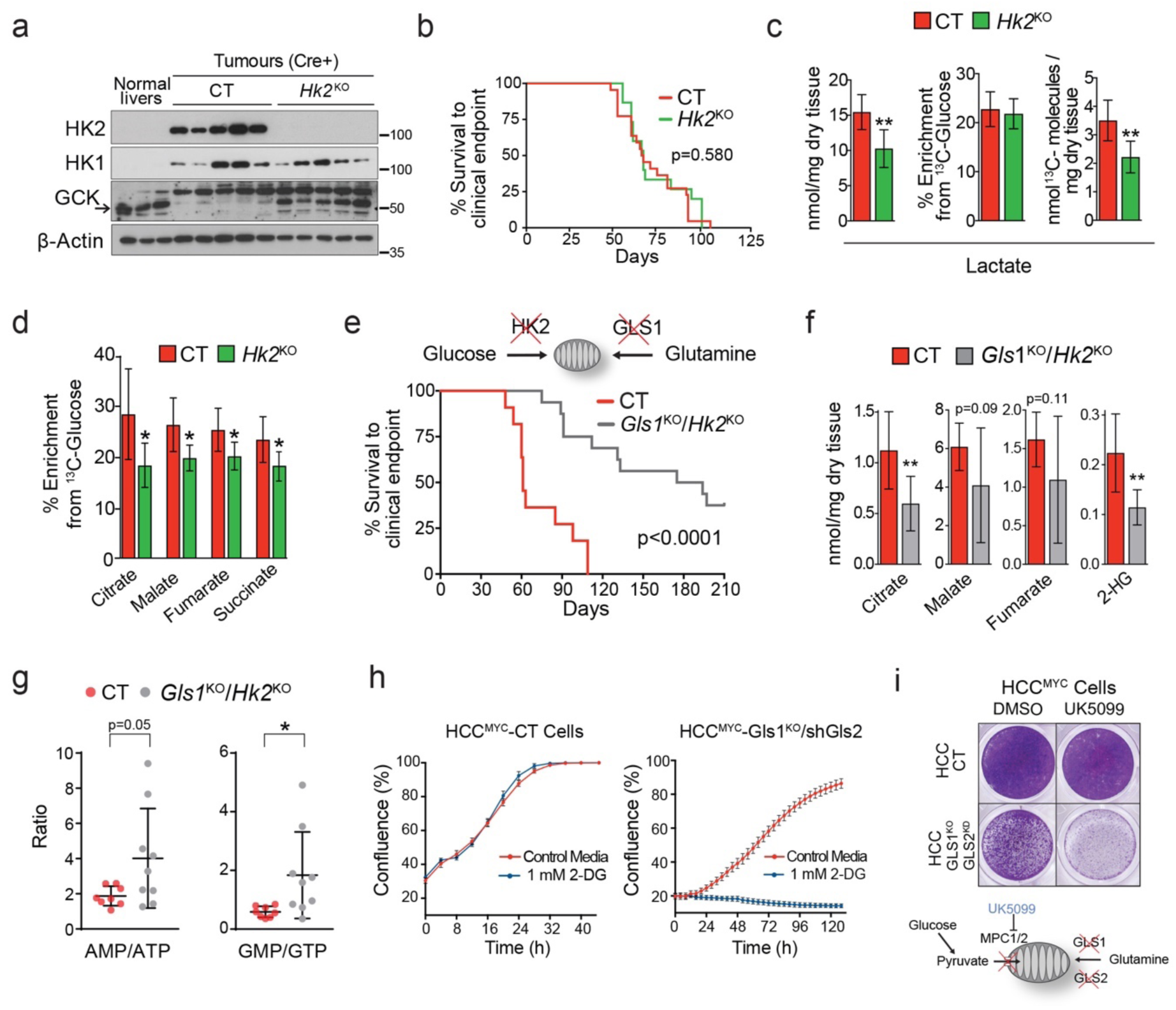
A cross-compensatory metabolism of glucose and glutamine sustains mitochondrial metabolic pools and MYC-induced tumourigenesis. (a-d) *Hk2* deletion in MYC-driven tumours decreases glycolysis but does not affect tumour burden. Tumours were induced by hydrodynamics-based transfection of MYC/MCL1 2 weeks after tamoxifen-induced liver-specific CRE activation in *Hk2*^fl/fl^/Alb-CreER^T2^/Rosa26eYFP and Alb-CreER^T2^/Rosa26eYFP mice. (a) Western blot demonstrating the absence of HK2 in *Hk2*^KO^ tumours. (b) Kaplan-Meier Survival Curve (CT n=21; *Hk2*^KO^ n=15). (c) Total content, ^13^C-enrichment, and total ^13^C content of lactate. (d) ^13^C-enrichment of the Krebs Cycle intermediates in CT and *Hk2*^KO^ tumours after a [U-^13^C]-glucose bolus (n=6 per group; GC-MS). (e-g) Deletion of both *Gls1* and *Hk2* affects MYC-induced tumourigenesis. Simultaneous Gls1 and Hk2 gene deletion and cell transformation by MYC were achieved by co-injection of pT3-CMV-Cre with c-MYC and MCL1-expressing plasmids. (e) Kaplan-Meier Survival Curve (CT n=11; *Gls1*^KO^/*Hk2*^KO^ n=16). (f) Total content of the Krebs cycle intermediates and 2-hydroxybutyrate (n= 8, 9, group labels as above, GC-MS). (g) AMP/ATP and GMP/GTP ratios (n= 8, 9, group labels as above, LC-MS); Data are presented as the mean ± S.D., *, *p* < 0.05; **, *p*< 0.01; ***, *p* < 0.001. (h) HCC^MYC^-CT and HCC^MYC^-*Gls1*^KO^/sh*Gls2* tumour cells were treated with either 0.5 or 1 mM 2-DG and growth was monitored in an IncuCyte Live-Cell analysis system. A representative of three independent experiments is shown. (i) HCC^MYC^-CT and HCC^MYC^-*Gls1*^KO^/sh*Gls2* tumour cells were treated with 100 µM UK5099 for 48 hours. Crystal violet staining from a representative of three independent experiments is shown. See also Extended data Fig 2 and 7

We next combined the deletion of both *Gls1* and *Hk2* (*Gls1*^KO^/*Hk2*^KO^) in MYC-induced liver tumours by co-injecting the MYC/MCL1 and CRE-expressing plasmids in *Gls1*^fl/fl^/*Hk*^fl/fl^, and wild type mice inducing the deletion of both *Gls1* and *HK2* at the tumour initiation stage. The deletion of both *Gls1* and *HK2* strikingly impaired tumour formation (Fig. 5e), with 37.5 % of mice not developing tumours even after one year. In contrast to either *Gls1*^KO^ or *Hk2*^KO^ tumours, *Gls1*^KO^/*Hk2*^KO^ tumours had lower levels of the Krebs cycle intermediates (Fig. 5f and Extended Data Fig. 7c). *Gls1*^KO^/*Hk2*^KO^ tumours had increased levels of glutamine consistent with the inhibition of glutamine catabolism (Extended Data Fig. 7d). However, they maintained the levels of lactate (Extended Data Fig. 7e) and amino acids (Extended Data Fig. 7f). Importantly, the AMP/ATP and GMP/GTP ratios were decreased in *Gls1*^KO^/*Hk2*^KO^ tumours in comparison with CT tumours (Fig. 5g) suggesting that *Gls1*^KO^/*Hk2*^KO^ tumours may fail to maintain a proper energy balance. Interestingly, *Gls1*^KO^/*Hk2*^KO^ tumours also had decreased levels of pentose phosphate pathway intermediates (Extended Data Fig. 7g). In general, the tumours that did develop showed a wide range of metabolite concentrations, consistent with a heterogeneous pattern of GLS2, GCK and HK1 expression (Extended Data Fig. 7h,i) suggesting different mechanisms of metabolic adaptation. Consistent with the previously published results^17^, some tumours also showed the residual expression of HK2. This result suggests that some of the tumours could have originated from cells with incomplete deletion of the gene and reinforces the idea that the deletion of both HK2 and Gls1 is highly detrimental for tumourigenesis. Overall, these results demonstrate that, when the catabolism of either glucose or glutamine into the Krebs cycle is inhibited, the corresponding metabolite can compensate to support the Krebs cycle activity and downstream pathways. Consistent with this concept, pharmacological inhibition of the mitochondrial pyruvate transporter, UK5099, or treatment with 2-Deoxy-D-Glucose (2DG) affected the proliferation of *Gls1*^KO^/sh*Gls2* cells more than of CT cells (Fig. 5h,i). These results suggest that reducing both anaplerotic pathways simultaneously could inhibit tumourigenesis more efficiently than inhibiting each pathway individually.

### Dietary interventions synergize with the inhibition of tumour biosynthetic pathways

One of the pathways that is fuelled by both glucose and glutamine in MYC-induced liver tumours is fatty acid biosynthesis (Fig. 1c,d). We tested whether this major pathway is required for MYC-induced tumourigenesis. To that effect we induced *Fasn* knockout at the onset of MYC-induced tumourigenesis (*Fasn*^fl/fl^+Alb-CreER^T2^+Tamoxifen; *Fasn*^KO^). *Fasn* KO delayed the onset of palpable tumours but did not increase the total time until a clinical endpoint (Fig. 6a,b). A total block of the incorporation of [U-^13^C]-glucose-derived carbons into fatty acids and unaffected labelling of citrate confirmed the inhibition of *de novo* lipogenesis in tumours at the FASN step (Fig. 6c,d and Extended Data Fig. 8a). Interestingly, the defect in lipogenesis in *Fasn*^KO^ tumours was not associated with the decrease in total content of fatty acids but rather with the change in the tumour fatty acid composition (lower c16:0/c18:0 and c16:0/c18:2 ratios) (Extended Data Fig. 8b,c). This outcome suggested that in the absence of *Fasn* tumour development was sustained by different exogenous lipids, and that in the absence of endogenous production, the lipid composition was determined by the lipid composition of the diet.

**Fig. 6.**
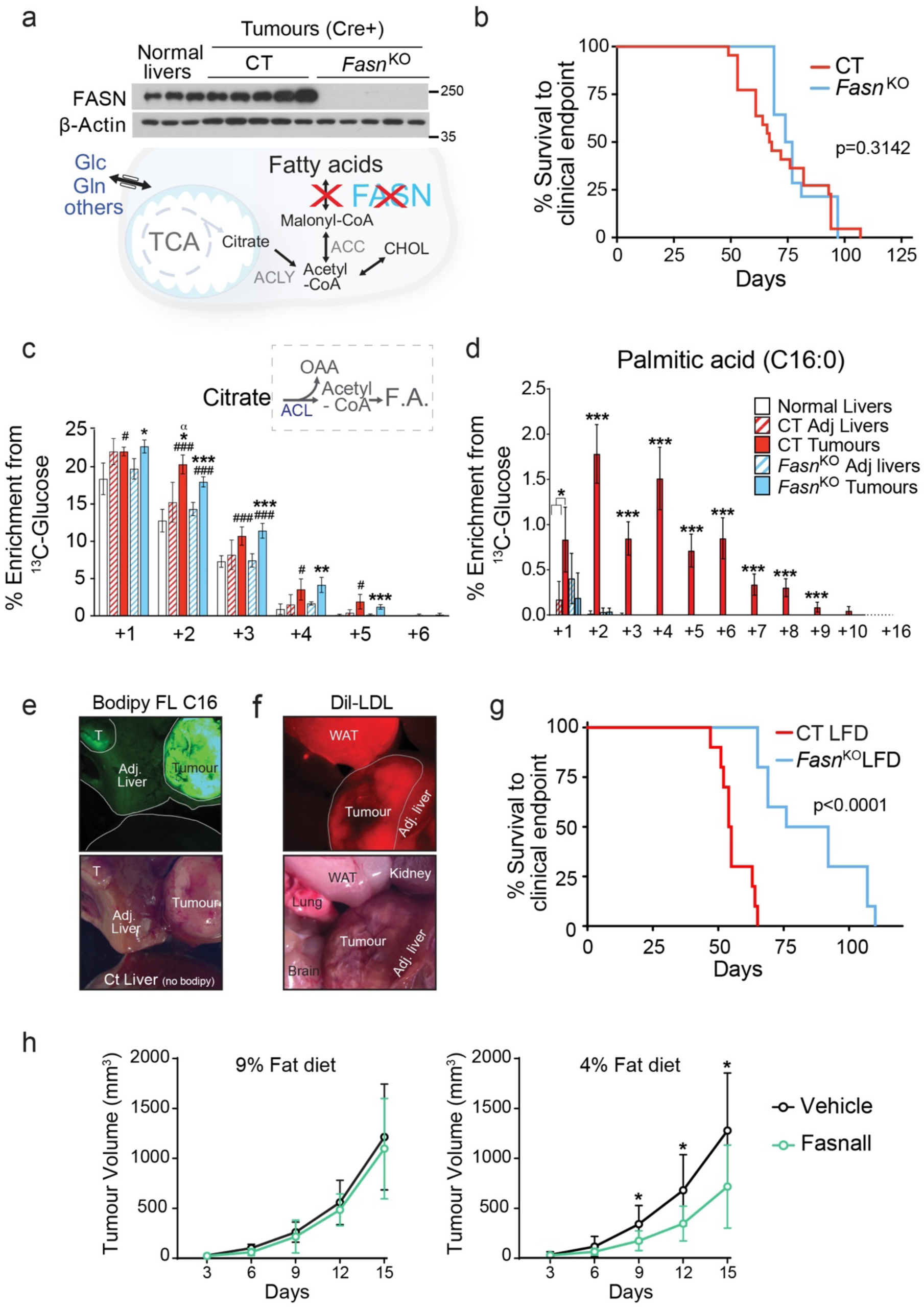
Lipid demands of tumours are fulfilled by a joint effort of *de novo* synthesis and uptake. (a-d) *Fasn* deletion in MYC-driven tumours ablates lipogenesis but does not significantly affect tumour burden. Tumours were induced by hydrodynamics-based transfection of MYC/MCL1 2 weeks after tamoxifen-induced liver-specific CRE activation in Fasn^fl/fl^/Alb-CreER^T2^/Rosa26eYFP and Alb-CreER^T2^/Rosa26eYFP mice. (a) Western blot demonstrating the absence of FASN in *Fasn*^KO^ tumours. (b) Kaplan-Meier Survival Curve (CT n=21; *Fasn*^KO^ n=14). (c and d), ^13^C-enrichment of citrate (n= 4,3,3,4,4; GC-MS) (c) and palmitic acid (n= 5,4,4,5,5; GC-MS) (d) in the indicated tissues after a [U-^13^C]-glucose infusion. (e and f) Direct fluorescence images of tissues of interest showing tumoural lipid uptake after *Fasn*^KO^ mice where intraperitoneally injected with fluorescently-labelled fatty acids (e) or fluorescently-labelled lipoproteins (f). (g) Kaplan-Meier Survival Curve of tumour bearing CT and *Fasn*^KO^ fed with the LFD (CT on the LFD n=8; *Fasn*^KO^ on the LFD n=10). Gene deletion was achieved during tumour induction by co-injection of pT3-CMV-Cre with c-MYC and MCL1-expressing plasmids in Fasn^fl/fl^ and WT mice. Animals were moved onto an indicated diet 1 week later. (h) The effect of combining the LFD and Fasn inhibitor, Fasnall, on the progression of MYC tumour cell-derived allografts. (Veh/9% fat diet n= 9; Fasnall/9% fat diet n= 9; Veh/4% fat diet n= 11; Fasnall/4% fat diet n= 11). Data are presented as the mean ± S.D. In (d) *, *p* < 0.05; **, *p*< 0.01; ***, *p* < 0.001, with respect to the rest of the groups, otherwise indicated. In (c) *, *vs* its own adjacent; # *vs* the normal liver; α, *vs Fasn*^KO^. See also Extended data Fig 2 and 8.

MYC-induced tumours express a wide range of lipid transporters (Extended Data Fig. 8d), suggesting that they may not rely on a single lipid uptake mechanism. Indeed, injecting a fluorescently-labelled free fatty acid (Bodipy™ FL C_16_) or fluorescently-labelled lipoproteins (Dil-LDL) demonstrated that both CT and *Fasn*^KO^ tumours imported serum lipids more avidly than the adjacent liver and most other tissues (Fig. 6e,f). We thus hypothesised that reducing lipid supply could synergize with *Fasn* deletion. To test whether modulating lipid availability *in vivo* affects tumour latency, we placed mice bearing *Fasn*^KO^ and CT tumours onto a diet with a lower fat content (LFD, 5.1% in comparison with 9% in our normal chow diet). We avoided the use of a zero-fat diet as it has been reported to induce a paradoxical lipid accumulation in *Fasn*^KO^ non-cancerous livers^36^. The LFD significantly increased the latency of *Fasn*^KO^ tumours (Fig. 6g), demonstrating that a sufficient level of circulating lipids was required for *Fasn*^KO^ tumours to grow at a rate comparable to CT tumours. Paradoxically, *Fasn*^KO^ tumours of mice on the LFD showed a tendency to increase lipid pools (Extended Data Fig. 8e), measured at the endpoint, but again with the change in the fatty acid composition, reflected by decreased 16:0/18:0 and 16:0/18:2 ratios (Extended Data Fig. 8f). Consistent with the *in vivo* results, cells isolated from *Fasn*^KO^ tumours showed a lower proliferation rate compared with CT cells in complete media and stopped proliferating in lipid-depleted media, which in both cases was rescued by fatty acid supplementation (Extended Data Fig. 8g).

To test whether diet fat content would also impact a therapeutic potential of pharmacological inhibition of FASN *in vivo* we used a previously-published Fasn inhibitor, Fasnall, which has shown an anti-tumour effect in mouse models of breast cancer^37^. HCC^MYC^ cells were injected into the flank of FVB/N mice and, once a tumour was first detected, mice were placed on either 9% or 4% fat diet and either Fasnall or vehicle was administered every 72 hours. Fig. 6h shows that a significant decrease in tumour growth was only observed when Fasnall was combined with the diet with lower fat content. Together, these results demonstrate that lipid requirements of tumours *in vivo* are sustained by both *de novo* lipogenesis and lipid uptake, and that strategies that do not target both pathways could be less efficient.

Another major pathway fuelled by both glucose and glutamine, shown to support the proliferation of cancer cells, is serine biosynthesis^38^. Serine and its immediate product glycine are essential precursors for protein, nucleic acid, folate, and lipid synthesis^38–41^. PSAT1 catalyzes one of the key steps in serine biosynthesis when glucose-derived carbon is combined with glutamine-derived nitrogen (Fig. 7a). MYC-induced liver tumours had increased serine biosynthetic capacity and significantly increased *Psat1* mRNA and protein levels (Fig. 1a,c,e; Extended Data Fig. 1b, 2b and Fig. 7b). Thus, we tested whether directly inhibiting serine biosynthesis would affect MYC-induced tumourigenesis. Surprisingly, *Psat1* deletion at the time of tumour initiation (*Psat1*^l/fl^+Alb-CreER^T2^+Tamoxifen; *Psat1*^KO^) had no effect on a tumour onset or progression (Fig. 7b,c). [U-^13^C]-glucose infusions and amino-^15^N-glutamine boluses demonstrated the ablation of *de novo* serine and glycine synthesis in *Psat1*^KO^ tumours (Fig. 7d, Extended Data Fig. 9a,b). Interestingly, only serine but not glycine levels were reduced (Fig. 7d), suggesting that serine pools were more dependent on serine synthesis, while glycine levels may be sustained by direct import from the blood stream. To test whether a reduction in serine/glycine (SG) availability *in vivo* would synergise with the absence of PSAT1, we placed mice with either *Psat1*^KO^ or CT tumours onto a SG-deficient diet (-SG) (Fig. 7e). The -SG diet has been shown to affect tumour progression in some mouse models of cancer^42^. Remarkably, the -SG diet had no significant effect on the latency of CT tumours (Fig. 7f), in which glycine, but not serine, levels were significantly reduced (Fig. 7g). Only the combination of *Psat1* deletion with the -SG diet achieved a reduction in the SG levels, and a striking delay in tumour progression (Fig. 7f). Consistent with the *in vivo* results, Psat1^WT^ but no Psat1^KO^ tumour cells were able to proliferate without serine/glycine and Psat1^KO^ cells were able to proliferate without glycine but not without serine. In the presence of glycine, Psat1^KO^ cells were only able to proliferate when formate, a carbon donor for one-carbon metabolism normally present in serum, was supplied (Extended Data Fig. 9c). These data demonstrate the need for targeting both the synthesis and the dietary supply of serine and glycine, and also suggests a limitation in the interconversion of these amino acids. Indeed, one of the factors preventing the serine production from glycine can be the depletion of methyl groups from one-carbon metabolism, which would limit nucleotide biosynthesis.

**Fig. 7.**
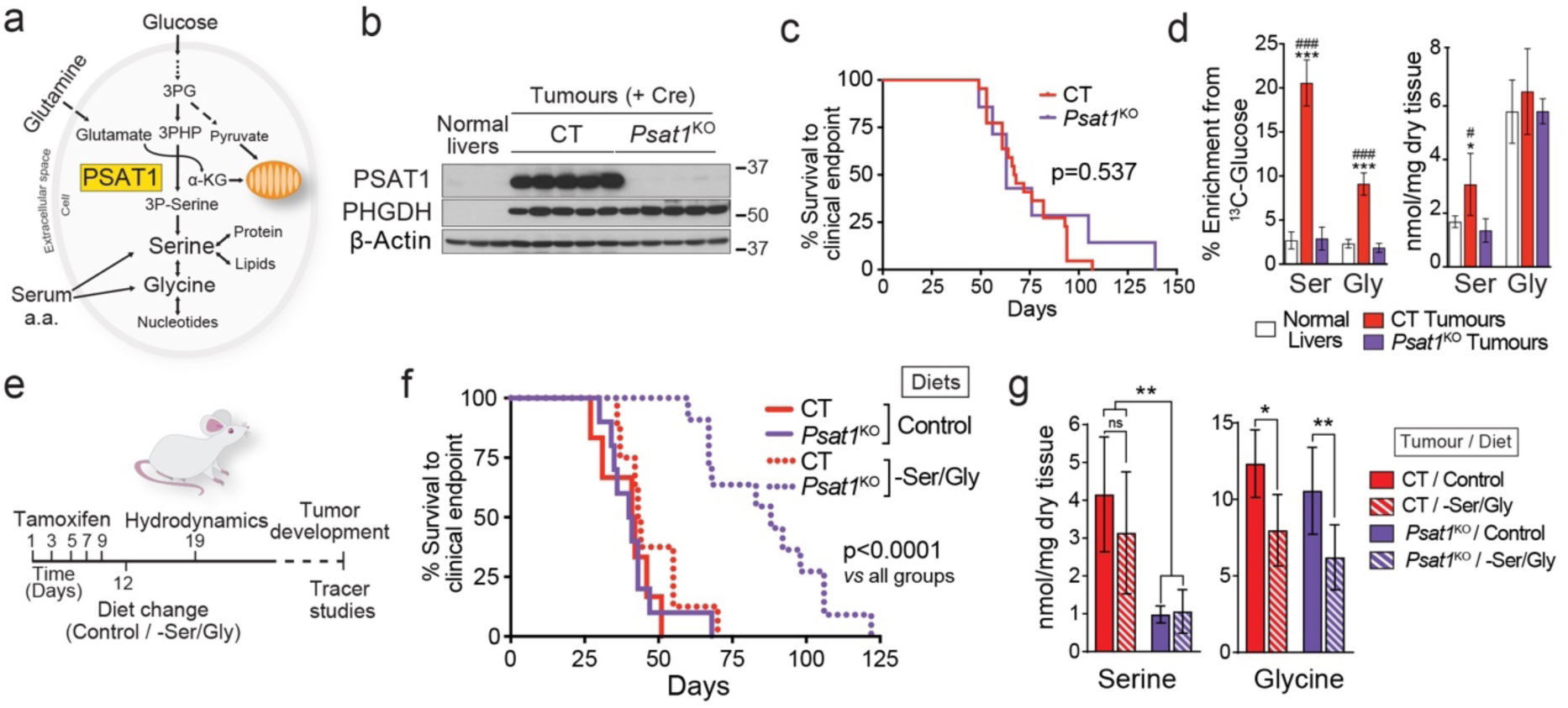
Depleting serine and glycine in tumours requires simultaneous intervention against endogenous production and a circulating supply. Psat1 KO was induced in hepatocytes of Psat1^fl/fl^/Alb-CreER^T2^/Rosa26eYFP mice by tamoxifen administration followed by hydrodynamics-based transfection of MYC/MCL1. (a) PSAT1 links glycolysis, glutaminolysis and TCA cycle. PSAT1 transfers the amino group from glutamate to the glucose derived P-pyruvate, and generates the Krebs cycle intermediate, α-ketoglutarate, and P-serine. (b) Western blot demonstrating the absence of PSAT1 in *Psat1*^KO^ tumours. (c) Kaplan-Meier Survival Curve (CT n=21; *Psat1*^KO^ n=7). (d) Total content, and ^13^C-enrichment of serine and glycine in the indicated tissues after a [U-^13^C]-glucose infusion (Normal livers n=6; CT n=5; *Psat1*^KO^ n=5; GC-MS). (e) Time scheme for the experiment combining *Psat1* knockout and serine and glycine deficient diet. (f) Kaplan-Meier Survival Curve (CT on the Control diet n=6; *Psat1*^KO^ on the Control diet n=10; CT on the –SG diet n=8; *Psat1*^KO^ on the –SG diet n=11). (g) Total content, and ^15^N-enrichment of serine and glycine in the indicated tumours after amino-^15^N-glutamine bolus in mice fed with either the Control or the -SG diets (n=5,5,7,7). Data are presented as the mean ± S.D., *, *p* < 0.05; **, *p*< 0.01; ***, *p* < 0.001; in D #, *p* < 0.05; ###, *p* < 0.001 relative to the normal liver, two-tailed t-test. See also Extended data Fig 2 and 9.

Surprisingly, a very profound delay in the formation of *Psat1*^KO^/–SG tumours was associated with only a few metabolic changes that we were able to detect. *Psat1*^KO^/–SG tumours had decreased levels of α-ketoglutarate but not any other Krebs cycle intermediates (Extended Data Fig. 9d). While Diehl F.F. et al^43^ reported a significant decrease in purine nucleotide biosynthesis in cancer cell lines deprived of serine, we did not observe any effect of -SG diet on the levels of purine nucleotides possibly due to 50% of serine still being present in serum of animals on -SG diet (Extended Data Fig. 9e). The appearance of ^15^N-serine and ^15^N-glycine in the blood of tumour-bearing animals injected with amino-^15^N-glutamine demonstrated that in the absence of dietary SG, blood SG levels can be sustained by synthesis and export from other organs (Extended Data Fig. 9e).

One of the differences that was observed, however, upon *Psat1^KO^* and -SG diet was in the levels of alanine, which was significantly increased in *Psat1^KO^/–SG* tumours (Extended Data Fig. 9a,d). Serine depletion has been shown to affect ROS balance^42^, and alter the metabolism of glycerophospholipids and sphingolipids^41^, leading to intracellular accumulation of cytotoxic deoxysphingolipids^44^ produced from alanine instead of serine. The latter can be one of the causes of the increased latency of *Psat1*^KO^/–SG tumours, which presented an increased alanine to serine ratio.

Together these data demonstrate that both SG synthesis and uptake can support the progression of MYC-induced liver tumours, and that a 50% reduction in the blood serine levels is sufficient to synergize with the inhibition of serine biosynthesis and profoundly impairs tumourigenesis.

## Discussion

A number of compounds that inhibit metabolic enzymes, including GLS1 and FASN, are now in different phases of clinical trials as potential cancer therapeutics. However metabolic flexibility could be one of the major obstacles to their success. Our *in vivo* results illustrate how cancer cells can withstand interventions that target metabolism, and explore different options for combined targeting of specific metabolic adaptations for greater impact on tumour development.

Our data builds up on the previous work that has shown promising but heterogeneous outcomes with the inhibition of individual metabolic enzymes in mouse models^17, 18, 20, 21, 45^. For example, although *Gls1* haplo-insufficiency (*Gls*^+/–^) slowed the early stage of tumour progression, and the treatment with the GLS1 inhibitor BPTES improved overall survival of mice with MYC-driven liver tumours^18^, lung tumours induced by KRas^G12D^ were not sensitive to pharmacological GLS1 inhibition and showed higher dependence on glucose catabolism through the Krebs cycle^4^. Consistently, *Hk2* deletion significantly inhibited the formation of KRas^LSL-G12D^-induced lung tumours^17^. *Hk2* deficiency also reduced the number of tumours in a diethylnitrosamine (DEN)-induced liver tumour model^19^. These studies contrasted with the absence of any effect of *Hk2* deletion in our model. Altogether, these data suggest that both the tissue of origin and the genetic drivers can not only influence the metabolic reprogramming during oncogenic transformation^10^ but also the mechanisms of metabolic adaptation.

Deleting the isoforms driving glucose and glutamine catabolism in a single tumour system allowed us to reveal several mechanisms of metabolic compensation that should be taken into account when designing metabolism-based therapeutic strategies. Inhibiting the expression of either *Hk2* or *Gls1* at the onset of MYC-induced liver tumourigenesis resulted in the formation of tumours with re-expression of *Gck* or *Gls2*, respectively, the isoforms expressed in the normal liver and repressed in tumours. We cannot exclude that the expression of liver isoforms in tumours is the result of the selection of transformed cells that maintain the expression of these isoforms during the transformation process. Nevertheless, these observations demonstrate that the expression of the enzyme isoforms in a parental tissue can determine the degree of tumour sensitivity to a metabolism-focused therapy. The mechanism that drives the expression of isoforms are yet to be elucidated. Furthermore, our results emphasize the role of glucose and glutamine as anaplerotic nutrients that support the Krebs cycle activity. The flux of carbons from each substrate can compensate and support the Krebs cycle intermediate pools when the other is limited. Indeed, only reducing both glucose and glutamine catabolism significantly reduced these pools and affected tumour formation. The results further substantiate previous elegant studies on the interactions between glucose and glutamine catabolism that support the survival of cancer cells^22–25^. This phenomenon is exemplified by the observed increase in glutamine catabolism upon inhibition of the mitochondrial pyruvate transporter^22^, or in response to a decreased rate of glycolysis when either mTOR or EGFR are targeted^24, 25^.

Furthermore, our results demonstrate that not only GLS1 but synergistic action of glutaminases and amidotransferases can fuel glutamine-derived carbons to the Krebs cycle. This is clinically-relevant since the expression of Gls1 and different amidotransferases strongly correlates in human HCCs suggesting that these patients may only benefit from a combination of compounds inhibiting these enzymes. Clinical trials testing DON in human tumours relied on the believe that this compound was also inhibiting glutaminases, based on previous results *in vitro* using relatively high doses^46, 47^. In contrast, our results suggest that DON-mediated inhibition of glutaminase cannot be achieved at therapeutically relevant doses. These findings open the door to exploring combinations of GLS1 inhibitors with inhibitors of other glutamine-catabolising enzymes or with compounds that suppress an activity of a glutamine-utilising pathway, such as DON. Importantly, DON prodrugs have been developed to enhance the delivery of an active compound to tumour tissues. These compounds circumvent the gastrointestinal toxicity and induce a dose-dependent inhibition of cell proliferation^48, 49^.

Finally, our results shed light on the key roles played by dietary intake and host tissues in supplying rescuing nutrients that allow tumour cells to shift to an auxotrophic phenotype, emphasizing the potential importance of dietary intervention as an adjuvant cancer therapy. Strategies that individually target serine synthesis, or reduce its availability, have been the focus of recent studies. The -SG diet affects tumour growth in some models, including Eµ-Myc lymphoma and Apc^min^-induced intestinal tumours while leaving KRas-induced pancreatic and intestinal tumours intact^42^. The -SG diet also enhanced the antineoplastic activity of biguadines on colon adenocarcinoma allografts, but had only a minor effect as a single therapy^50^. Our results suggest that the ability of tumours to synthesise serine makes them resistant to the -SG diet; inhibiting serine biosynthesis is necessary to broaden the spectrum of cancer types that could respond positively to this dietary regime. Importantly, inhibitors of serine biosynthesis pathway have been developed^39, 51^, and the sensitivity towards these inhibitors may also be determined by the concentration of serine and glycine in a specific tissue^52^. Together with inhibiting SG biosynthesis in tumours, these compounds should also be able to suppress the synthesis of SG by the host tissues. In this context a partial reduction of dietary serine and glycine, which is more feasible to implement in a clinic than a total depletion, might be ample to deplete serine/glycine serum levels and affect tumour progression.

Similar to the situation with serine biosynthesis, different tumours have varying requirements for *de novo* fatty acid synthesis^21, 45^. FASN inhibitor TVB2640, alone or in combination, is currently being tested in different clinical trials (ClinicalTrials.gov identifiers: NCT02223247, NCT03032484, NCT02980029, and NCT03179904), and preliminarily shows anti-tumour activity^53^. Although low fat diets on their own seem to have minor effects on human cancer progression^54^, our results suggest that decreasing lipid availability or inhibiting lipid uptake should be considered as an adjuvant for therapies that target *de novo* lipogenesis. Although the inhibition of lipid transporters could be one option^55^, the presence of several uptake mechanisms could become a major obstacle in some tumour types, hence low-fat diets represent an attractive alternative.

In conclusion, our study illuminates how the understanding of metabolic adaptations *in vivo* is crucial for the design of effective therapeutic strategies for cancer, and that exploitation of combinatorial interventions against compensatory metabolic pathways may lead to more robust inhibition of tumour growth.

## Author Contributions

A.M.L. and M.Y. conceived the research project, designed and performed experiments and wrote the paper. W.L, P.C.D., N.L., J.I.M, L.N.V. performed research and/or contributed to analysis and discussion. S.R., N.H provided mouse models. V.L. and M.R.J provided human and mouse organoids. M.C., Z.W and N.P.J. provided Compound 27. X.C. and CX provided tissue microarray analysis, and reagents.

## Acknowledgements

This work was supported by the Francis Crick Institute which receives its core funding from Cancer Research UK (FC001223), the UK Medical Research Council (FC001223), and the Wellcome Trust (FC001223). This work was supported by grants from NIH R01CA136606 and R21CA198490 to XC; P30DK026743 for UCSF Liver Center; NIH R01CA206167 and Department of Veteran Affairs BX000733 for NX. We thank The Francis Crick Institute Biological Services for breeding and maintenance of the mice. We also thank Alex P. Gould, Patricia M. Nunes, and Andrew P. Bailey for critical reading and useful comments on the manuscript.

## Declaration of Interests

The authors declare no competing interests.

## Methods

### Mice

Mice transgenic for both LAP-tTA and tet-o-MYC that overexpress MYC in the liver were generated as previously^9^. To initiate MYC expression, mice were weaned into regular chow. Mice kept on a doxycycline-containing diet did not overexpress c-MYC and were used as controls (normal livers). *Gls1*^fl/fl^ mice^56^, *Hk2*^fl/fl^ mice^17^ and *Fasn^fl/fl^* mice^45^ were previously described. *Psat1*^fl/fl^ mice were derived from *Psat1*^tm1a(KOMP)Wtsi^ mice, obtained from the KOMP repository (UCDavis, MGI:4363603). *Gls1*^fl/fl^, *Hk2*^fl/fl^, *Psat1*^fl/fl^, and *Fasn*^fl/fl^ mice, originated in different backgrounds, were back-crossed for a minimum of 10 generations with an FVBN/J strain to allow comparison among all the experimental groups. All lines were crossed with Alb-CreER^T257^ and Rosa26eYFP lines to generate Gene^fl/fl^/Alb-CreER^T2^/Rosa26eYFP lines. Gene deletion was achieved by intraperitoneally (i.p.) administration of tamoxifen (Sigma; 10 mg/kg; dissolved in 1:10 ethanol:oil solution) starting at 5/6 weeks of age. Mice were bred and maintained under specific pathogen-free conditions at The Francis Crick Institute (Mill Hill laboratory and Midland Road Laboratory), and all procedures and animal husbandry were carried out in accordance with the UK Home Office, under the Animals (Scientific Procedures) Act 1986, and the Local Ethics Committee under the Project license number P609116C5.

### Constructs and reagents

The constructs for mouse injection, including pT3-EF1α-c-MYC, pT3-EF1α-MCL1 and pCMV-SB (which encodes for sleeping beauty transposase), and pCMV-CRE (which encodes for CRE recombinase), were previously described^58^. miR30 based shRNA targeting for *Gls2* and Renilla Luciferase were cloned into pT3-EF1α-c-MYC. The shRNA sequences are: *Gls2*: CATCATGCCAACAAGCAACTT and Renilla Luciferase: AGGAATTATAATGCTTATCTA. All plasmids used for *in vivo* experiments were purified using the Endotoxin-free Maxiprep kit (Qiagen). [U-^13^C]-glucose, [U-^13^C]-glutamine, ^15^N-glutamine and ^15^N-alanine were purchased from Cambridge Isotope Laboratories, Inc.

### Hydrodynamics-based transfection of DNA in the liver and mouse monitoring

Hydrodynamic-based transfection was performed as described^59^, with some variations. Briefly, 10µg of pT3-EF1α-c-MYC, pT3-EF1α-c-MYC/sh*Gls2* or pT3-EF1α-c-MYC/shLuc with 10 µg of pT3-EF1α-Mcl1 along with sleeping beauty transposase (SB) in a ratio of 25:1 were diluted in a volume of saline (0.9% NaCl) corresponding to 10% of body weight, and injected into the lateral tail vein of 7 to 9-week-old FVB/N mice in 7 to 9 seconds. Where indicated, gene deletion was achieved by co-injection of 40 µg of CRE-recombinase (pT3-CMV-Cre) with c-MYC and MCL1-expressing plasmids and 4 µg of SB-encoding plasmid. Titration of the amount of SB allows to ensure a reliable amount of integration events and induction of tumourigenesis. Mice used as controls (normal livers - Figures 2-7) where injected with an empty plasmid (pT3-EF1α-MCS). The timescale for tumour burden is specific for different methods of c-MYC induction and gene knockout.

### Xenograft and allograft experiments

5 × 10^6^ human HCT116 cells were subcutaneously injected into the flank of 6-8-week-old SCID mice. Xenografts were allowed to grow 2-3 mm before randomizing the mice into groups. Tumours were measured with caliper every 3 days and the volume is calculated with the formula. All tumours were harvested at the end of the experiment and stored for further processing.

### Diets and compound administration

For the dietary amino acid restriction two synthetic diets were used: TestDiet® Baker amino acid diet with no added serine and glycine 5W53, and the control diet Baker amino acid defined diet 5CC7. Mice were placed on diets one week before hydrodynamics. For the low-fat diet experiment a Low-Fat Control for Western Diet (5TJS) from Test Diet was used. To improve acceptance of the diet, in this experiment 5TJS was initially mixed with 5CC7 pellets (both diets have 5.1% fat content). Mice were placed on diets one week after hydrodynamics. For the treatment with the Fasn inhibitor Fasnall, a 4% and 9% Fat diets from Teklad Global Diets were used (2914C and 2919 respectively). Diet was changed 6 days after tumour injection, when tumours reached 0.3 mm of diametre, and 25mg/kg of Fasnall was administered every 72 hours i.p from 1 day after diet change. The vehicle was 50% DMSO, 25% saline, 25% PBS. For short-term DON administration, 50 mg/kg of DON was administered 4 hours before labelling. For long terms treatment. For the treatment of mice bearing HCT116 xenografts 25 mg/kg of DON and 100mg/kg of Gls1 inhibitor (compound 27) were used. DON was dissolved in saline and administered every 72 hours via intraperitoneal. Gls1i/Compound 27^35^ was dissolved at 10 mg/ml in 1% tween 80 and the pH was adjusted to 3.5, and kept at room temperature under constant mixing for a maximum of 7 days. Compound 27 (and vehicle) were administered every 24 hours via oral gavage.

### Stable isotope labelling *in vivo*

Two types of *in vivo* label administration were performed: either bolus injections (once for glucose – endpoint: 15 min; twice for glutamine– endpoint: 30 min; and one for alanine - endpoint: 30 min), or a long-term infusion (3 hours). Boluses of stable isotope-labelled compounds dissolved in saline (0.9% NaCl) were administered through the tail-vein (i.v.). [U-^13^C]-glucose was administered in a single bolus of 570 mg/kg (calculated as 20 mg of [U-^13^C]-glucose for an average body weight (b.w.) of 35g dissolved in 100 µl of saline). Due to a limited solubility, administration of glutamine-derived stable isotopes ([U-^13^C]-glutamine, amino-^15^N-glutamine and amido-^15^N-glutamine) was divided into 2 boluses separated by a 15-minute interval. A total dose of glutamine-derived stable isotopes was 340 mg/kg (calculated as 6+6 mg of [U-^13^C]-glutamine for an average body weight of 35g).^15^N-alanine was administered in a single bolus of 250 mg/kg.

Infusions were performed in animals under isofluorane anaesthesia, through a tail vein catheter, using an Aladdin AL-1000 pump (World Precision Instruments). The infusions protocol was based on the experiments described previously^60^. For [U-^13^C]-glucose, mice received a 400 mg/kg bolus, followed by a 12 µg/kg of b.w./hour infusion for 3 hours at 0.15 ml/hour. For glutamine-derived stable isotopes infusions, mice received a 187 mg/kg bolus, followed for a 5 µg/Kg of b.w./hour infusion for 3 hours at 0.15 ml/hour. At the endpoint blood was obtained by cardiac puncture under terminal anaesthesia, and tissues were then harvested and freeze-clamped with a Wollenberger-like device precooled in liquid nitrogen. The tissues were stored at −80°C until analysis.

### Tissue metabolite extraction and analysis

Freeze-clamped tissues were ground in liquid nitrogen with mortar and pestle and subsequently lyophilized on a FreeZone 4.5 Freeze Dry System (Labconco). Metabolite extraction from mouse tissues and derivatization of polar samples were performed as previously described^61^. Briefly, 10-15 mg of dry weight per sample were extracted with 1.8 ml of chloroform:methanol (2:1 v/v) containing a combinations of internal standards for 1 hour at 4°C with intermittent sonication. Standards include scylloinositol (10 nmol), L-Norleucine (10 nmol), ^13^C_5_-^15^N-Valine (5 µM final), 4,4-dimethyl-4-silapentane-1-sulfonic acid (DSS, 1 mM final), added with the methanol, and margaric acid (c17:0, 10-20 µg) added in the chloroform) After centrifugation (18,000g for 10 minutes at 4°C), the supernatant (SN1) was vacuum dried in rotational-vacuum-concentrator RVC 2-33 CD (Christ). The pellet was re-extracted with methanol:water (2:1 v/v) as described above. After centrifugation, supernatant (SN2) was vacuum dried in the SN1 tube. Phase partitioning (chloroform:methanol:water, 1:3:3 v/v) was used to separate polar and apolar metabolites. Fractions were vacuum dried as described above.

For GC-MS analysis of the polar metabolites, a part of the polar fraction was washed twice with methanol, derivatized by methoximation (Sigma, 20 µl, 20 mg/ml in pyridine) and trimethylsilylation (20 µl of N,O-bis(trimethylsilyl)trifluoroacetamide reagent (BSTFA) containing 1% trimethylchlorosilane (TMCS), Supelco), and analysed on an Agilent 7890A-5975C GC-MS system^61, 62^. Splitless injection (injection temperature 270°C) onto a 30 m + 10 m × 0.25 mm DB-5MS+DG column (Agilent J&W) was used, using helium as the carrier gas, in electron ionization (EI) mode. The initial oven temperature was 70°C (2 min), followed by temperature gradients to 295°C at 12.5°C/min and then to 320°C 25°C/min (held for 3 min).

For GC-MS analysis of fatty acids, part of the apolar fraction was washed twice with methanol, derivatized by methoximation (Sigma, 20 µl, 20 mg/ml in pyridine) and analysed on an Agilent 7890A-5975C GC-MS system^61, 62^. Splitless injection (injection temperature 270°C) onto a 30 m + 10 m × 0.25 mm DB-5MS+DG column (Agilent J&W) was used, using helium as the carrier gas, in electron ionization (EI) mode. The initial oven temperature was 50°C (1 min), followed by temperature gradients to 190°C at 20°C/min (held for 3 min), then to 242°C 4°C/min, then to 292°C at 10°C/min, and then to 320°C at 20°C/min. Triglycerides were purified by solid phase extraction on aminopropyl silica columns (Biotage Insolute NH2), with tripentadecanoin as an internal standard.

Metabolite quantification and isotopologue distributions were corrected for the occurrence of natural isotopes in both the metabolite and the derivatization reagent. Data analysis and peak quantifications were performed using MassHunter Quantitative Analysis software (B.06.00 SP01, Agilent Technologies). The level of labeling of individual metabolites was corrected for natural abundance of isotopes in both the metabolite and the derivatization reagent^58^. Abundance was calculated by comparison to responses of known amounts of authentic standards.

NMR spectra were acquired at 25°C with a Bruker Avance III HD instrument with a nominal ^1^H frequency of either 700 or 800 MHz using 3 mm tubes in a 5 mm CPTCI cryoprobe. For ^1^H 1D profiling spectra the Bruker pulse program *noesygppr1d* was used with a 1 s presaturation pulse (50 Hz bandwidth) centred on the water resonance, 0.1 ms mixing time, and 4 s acquisition time at 25°C. Typically 128 transients were acquired. To monitor the fate of ^15^N atoms derived from ^15^N-glutamine labelling we acquired 2D ^15^N,^1^H heteronuclear multiple bond correlation (HMBC) spectra at ^1^H 700 MHz using the Bruker pulse program *hmbcf3gpndqf*, adapted to include solvent water resonance irradiation during the relaxation delay. Typically, the acquisition parameters employed were sweep widths 13 ppm (^1^H) and 220 ppm (^15^N), with offsets on the solvent water signal (^1^H) and at 120 ppm (^15^N). Acquisition times were 0.86 s (^1^H) and 0.012 s (^15^N), and pulse widths 7.4 µs (^1^H) and 25 µs (^15^N). 16 transients were collected for each increment, yielding a total measurement time of 4 h 16 m. The spectra were processed and plotted using Bruker TopSpin 3.5. The raw data were apodized with 1 Hz line broadening in the ^1^H dimension and unshifted sine bell in the ^15^N dimension and zero-filled to a matrix size of 32K × 512 points. The spectra are presented in a magnitude mode. Carbon-13 incorporation was assessed using 2D ^13^C,^1^H-heteronuclear single quantum coherence (HSQC) spectroscopy with the pulse sequence *hsqcetgpsisp2* using sweep widths 14 ppm (^1^H) and 165 ppm (^13^C) and offsets on the solvent water signal (^1^H) and 70 ppm (^13^C). Acquisition times were 0.16 s (^1^H) and 0.0176 s (^13^C), and pulse widths 7.4 µs (^1^H) and 11 µs (^13^C). Eight transients were collected for each increment. Non-uniform sensing (35%) of the data points in the indirect dimension was employed, yielding a total measurement time of 1 h 46 m. The spectra were reconstructed using the compressed sensing algorithm in Topspin, using Lorentz-to-Gaussian transformation (LB = −1 Hz; GB = 0.08) and cosine apodization in the ^1^H and ^13^C dimensions, respectively, and zero-filling to 4K ′ 1K points. NMR spectra were analysed with rNMR software^63^.

Metabolite analysis was performed by LC-MS using a Q-EXACTIVE Plus (Orbitrap) mass spectrometer from Thermo Fisher Scientific (Bremen, Germany) coupled with a Vanquish UHPLC system from Thermo Fisher Scientific (Bremen, Germany). The chromatographic separation was performed on a SeQuant® Zic®pHILIC (Merck Millipore) column (5 µm particle size, polymeric, 150 × 4.6 mm). The injection volume was 10 µl, the oven temperature was maintained at 25°C, and the autosampler tray temperature was maintained at 4°C. Chromatographic separation was achieved using a gradient program at a constant flow rate of 300 µl/min over a total run time of 25 min. The elution gradient was programmed as decreasing percentage of B from 80% to 5% during 17 minutes, holding at 5% of B during 3 minutes and finally re-equilibrating the column at 80% of B during 4 minutes. Solvent A was 20 mM ammonium carbonate and solvent B was acetonitrile. Metabolites were identified and quantified by accurate mass and retention time and by comparison to the retention times, mass spectra, and responses of known amounts of authentic standards using TraceFinder 4.1 EFS software (Thermo Fisher Scientific). Label incorporation and abundance was estimated using TraceFinder 4.1 EFS software. The level of labelling of individual metabolites was estimated as the percentage of the metabolite pool containing one or more ^13^C atoms after correction for natural abundance isotopes. Abundance was given relatively to the internal standard.

### Cell experiments

For isolation of tumour cells, harvested tissue was maintained in the ice-cold serum-free DMEM until processing (not more than 10 minutes). Tumours were minced in a 10 cm Petri dish using a scalpel, and washed twice with ice-cold HBSS containing EGTA. Then, intensive additional mincing was performed and 10 ml of digestion medium (HBSS containing 4 mM Ca^2+^, 5.5 mM glucose, 2 mM glutamine, and 40 µg/ml of Liberase TM (Roche)) were added, and incubated for 15-20 minutes at 37°C. The digested mixture was then passed through a 100-micron sterile nylon mesh cell strainer into a sterile 50 ml conical tube. 100-micron mesh allowed small groups of cells to pass-through, which lead to better survival results in some genotypes. Filtered cells were then centrifuged for 5 min at 200×g. The supernatant was removed and cells were resuspended in the washing media (MEM-Eeagle for suspension culture plus 2 mM glutamine; Biological industries). Cells were then seeded in DMEM containing 25 mM glucose, 2 mM glutamine, 10% FBS, 1% Penicillin/Streptomicin. To knock-out *Psat1* in cells isolated from MYC-driven tumours induced in *Psat1*^fl/fl^ mice the retroviral vector MSCV-CreERT2 puro (Addgene plasmid #22776) was used. After transduction, cells were selected with puromycin, and then treated with 4-hydroxytamoxifen (4-OHT) to induce Cre activity. *Gls1*^KO^/sh*Luc, Gls1*^KO^/sh*Gls2* cells, *Fasn*^KO^ cells and their CT counterparts were directly isolated from the respective tumours. The experiments for the analysis by GC-MS were performed in medium containing 10 mM glucose and 2 mM glutamine in 6-well plates and 10 cm plates for NMR analysis. Metabolite extraction from cells was performed as described for tissues except the first step when the media was aspirated and cells were rapidly washed with ice-cold PBS and 600 µl of ice-cold methanol containing standards were added. Cells were scraped and transferred to an Eppendorf where 1.2 ml of chloroform were added. Cell proliferation was monitored by using the IncuCyte® system (Essen Bioscience). In each experiment, each condition was plated in triplicate. Growth curves were generated from data points acquired during 4 h interval imaging.

### Western blot and immunofluorescence

For western blotting tissues were homogenized in radioimmunoprecipitation assay buffer (RIPA) supplemented with protease and phosphatase inhibitor cocktails (aprotinin, leupeptin, benzamidin, pepstatin A, PMSF, sodium fluoride, sodium orthovanadate and β-glycerophosphate), with an Ultra-Turrax® homogenizer. Samples were centrifuged at 15,000g for 15 min at 4°C. Supernatantprotein was quantified and protein extracts were mixed with a loading buffer 4X (40% glycerol, 2% SDS, 250 mM Tris pH 6.8, 0.02% bromophenol blue, and 12% β-mercaptoethanol). Western blots were performed with 20 µg of tissue extracts. Proteins were separated in 8–12% SDS-PAGE and transferred to a nitrocellulose membrane (BioRad). The membrane was blocked in 5% milk, incubated with primary antibody in TBST with 5% BSA overnight at 4°C, and then incubated with Horseradish peroxidase activity linked to secondary antibody. Protein signal was detected with ECL substrate (Pierce) using X-Ray films. Primary antibodies used were c-MYC (1472-1, Epitomics), PSAT1 (20180-1-AP, Protein Tech), PHGDH (sc292792, Santa Cruz), β-Actin (A2228 and A3854, Sigma), Cleaved CASP3 (9661, Asp175, Cell Signaling), Glucokinase (AP7901C, Abgent), GLS2 (6217, ProSci), FASN (#23180, C20G5, Cell Signaling), HK1 (#2024, C35C4, Cell Signaling), HK2 (#2867, C64G5, Cell Signaling), PCNA (#13110, D3H8P, Cell Signaling), PARP (#9542, Cell Signaling), GLS1-KGA specific antibody was a generous gift from Astra Zeneca.

For immunofluorescence, tissues were fixed for 24 hours in 4% paraformaldehyde, equilibrated in 30% sucrose, embedded in OCT compound (VWR international), and stored at −80°C until 7-µm-thick cryosections were obtained. Sections were permeabilized by incubating with 0.2% Triton X-100 in PBS for 10 min, and treated with the indicated primary antibody for 16 hours. Primary antibodies used were: GFP (4745-1051, Bio-Rad) and PanCK (Z0622, Dako). After four washes with 0.05% Tween in PBS, cells were incubated with anti-rabbit Alexa Fluor® 488 or 555 secondary antibodies (Invitrogen) for 1 hour. Nuclear marker DAPI was included in the mounting media. Samples were examined using a confocal microscope (TCS SP5 II Leica) and LCS Lite software (Leica) was used to collect digital images.

### Mouse intestinal organoid culture

Organoids were stablished from tumours isolated from APC^min^ mice using a previously described protocol^64^, with an additional collagenase and dispase digestion step after the EDTA-chelation step^65^ where Matrigel was replaced with Cultrex® BME, Type 2 RGF PathClear (Amsbio 3533-010-02). Organoids were cultured in Intesticult^TM^ Organoid Growth media (06005, Stem Cell Technologies). The Rho kinase inhibitor Y-27632 (Sigma) was added to the culture when trypsinised.

### Human material and patient-derived organoid culture

Samples have been harvested during surgeries at the University College London Hospital, in accordance with ethical approval, REC Ref: 15/YH/0311. Written informed consent was obtained. Intestinal samples were obtained from patients with colorectal cancer. Crypts were isolated from human intestinal tissue by incubating for 1 hour with chelation buffer (5.6 mM Na_2_HPO_4_, 8 mM KH_2_PO_4_, 96 mM NaCl, 1.6 mM KCl, 44 mM sucrose, 54.8 mM D-sorbitol, 0.5 M EDTA and 1M DTT at 4°C, and plated in drops of BME (Sato et al, 2009). After polymerization culture media was added. Human intestinal organoid media contains advanced DMEM/F12 medium (Invitrogen) including B27 (Invitrogen), nicotinamide (Sigma-Aldrich), N-acetylcysteine (Sigma-Aldrich), EGF (Invitrogen PMG8043), TGF-ß type I receptor inhibitor A83-01 (Tocris), P38 inhibitor SB202190 (Sigma-Aldrich), gastrin I (Sigma-Aldrich), Wnt3a conditioned media (50% produced using stably transfected Lcells), Noggin and R-spondin conditioned media.

### Cell Titer Glo

Organoids were trypsinized using TrypLE and filtered with a 20 µm cell strainer. 3500 single cells were seeded in BME per 48 well and placed in a 37°C incubator to polymerize for 20 minutes. 250 µl of Intesticult^TM^ Organoid Growth media (06005, Stem Cell Technologies) (for APC^min^ organoids) or 250 µl of complete human organoid media (for human patient organoids) with the indicated treatments was added for the indicated times. The media were supplemented with 10 µM Y-27632 for two days after plating. Cell titer Glo Luminiscent Cell viability assay (G7572, Promega) was used to assess viability of the organoids. Experiments were performed at least three times with three triplicates each.

### Organoid colony formation assay

Organoids were dissociated into single cells using TrypLE and further filtered with a 20 µm cell strainer. For each condition, 3500 single cells were seeded in BME per 48 well and placed in a 37°C incubator to polymerize for 20 minutes. 250 µl of Intesticult^TM^ Organoid Growth media (06005, Stem Cell Technologies) (for APC^min^ organoids) or 250 µl of complete human organoid media (for human patient organoids) with the indicated conditions. The media were supplemented with 10 µM Y-27632 for two days after plating. Number of spheres formed in each well was counted as plating efficiency. Images were captured at the indicated times with EVOS FL cell imaging system (Life technologies). Experiments were performed at least three times with three triplicates each.

### Microarray analysis of mouse tissues

The microarray data from 4 MYC-induced liver tumours and 4 wild type control livers can be found on Gene Expression Omnibus (GEO) accession GSE129013 (http://www.ncbi.nlm.nih.gov/geo). Data were analysed using Bioconductor 2.13 running on R 3.0.2. Probeset expression measures were calculated using Affymetrix package’s Robust Multichip Average (RMA) default method.

### Statistical analysis

All analyses were performed using GraphPad Prism 7 software. Statistical analyses were performed applying a two-tailed Student’s t-test, One-Way ANOVA or Two-Way ANOVA, where indicated. Error bars in figures represent the standard deviation (S.D.). No statistical methods were used to predetermine sample size. When possible (for example, for shRNA experiments, or diets), mice were randomized into groups. Samples were randomized for metabolomics analysis. The investigators were not blinded to allocation during experiments.

### Data availability

All the data that support the findings of this study are available from the corresponding author on reasonable request.

## Supplemental Information Titles and Legends

**Extended Data Fig. 1.**
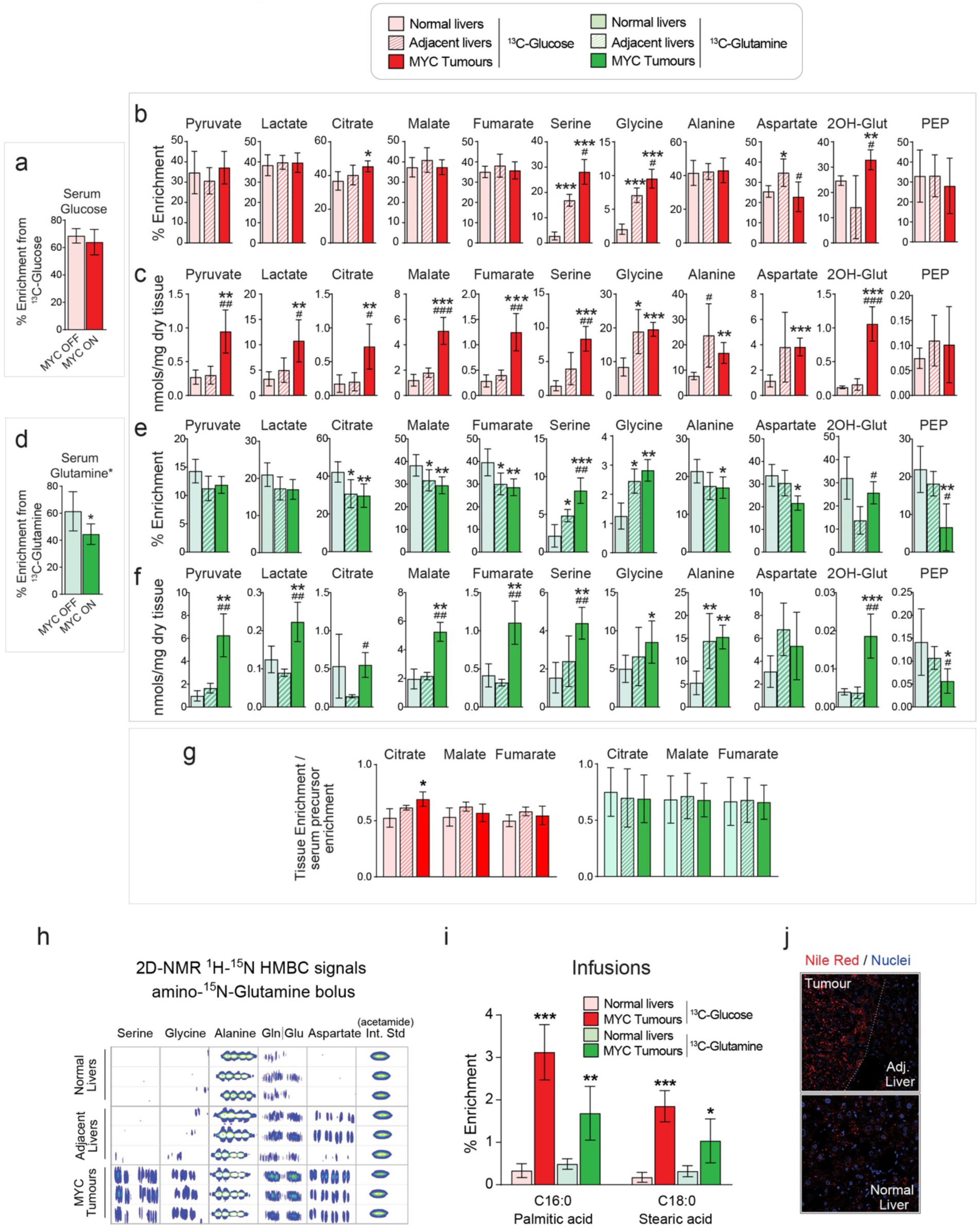
Glucose and glutamine metabolism in MYC liver tumours. Related to Fig. 1. (a-g) Mice bearing MYC-driven liver tumours and control mice (n = 5 per group) were infused with [U-^13^C]-glucose (a-c), or [U-^13^C]-glutamine (d-f) and the label incorporation into tissue metabolites was analysed by GC-MS: Percent enrichment in either serum glucose (a) or glutamine (d) in mice administered either [U-^13^C]-glucose or [U-^13^C]-glutamine bolus, respectively (*glutamine enrichment is estimated from quantification of its spontaneous product pyroglutamate). (b and e) Percent enrichment. (c and g) Total content of metabolites. Note that lower glutamine enrichment in the Krebs cycle intermediates in tumours in comparison with normal livers is proportional to the difference in serum enrichment between control and tumour-bearing mice; normalized values for the TCA cycle metabolites from (b and e) are shown in (g). Data are presented as the mean ± S.D., *, *p* < 0.05; **, *p*< 0.01; ***, *p* < 0.001, relative to normal livers; #, *p* < 0.05; ##, *p* < 0.01; ###, *p* < 0.001, relative to adjacent livers, two-tailed t-test. (h) ^15^N-HMBC 2D NMR signals of the indicated metabolites in the indicated mouse tissues after amino-^15^N-glutamine bolus. (i) Percent enrichment of tissue total fatty acids (both free and esterified) after either [U-^13^C]-glucose or [U-^13^C]-glutamine infusions (n=5 per group; GC-MS). (j) Nile red fluorescence visualised by confocal microscopy showing neutral lipid accumulation (in red) in MYC-driven tumours. Nuclei are shown in blue.

**Extended Data Fig. 2.**
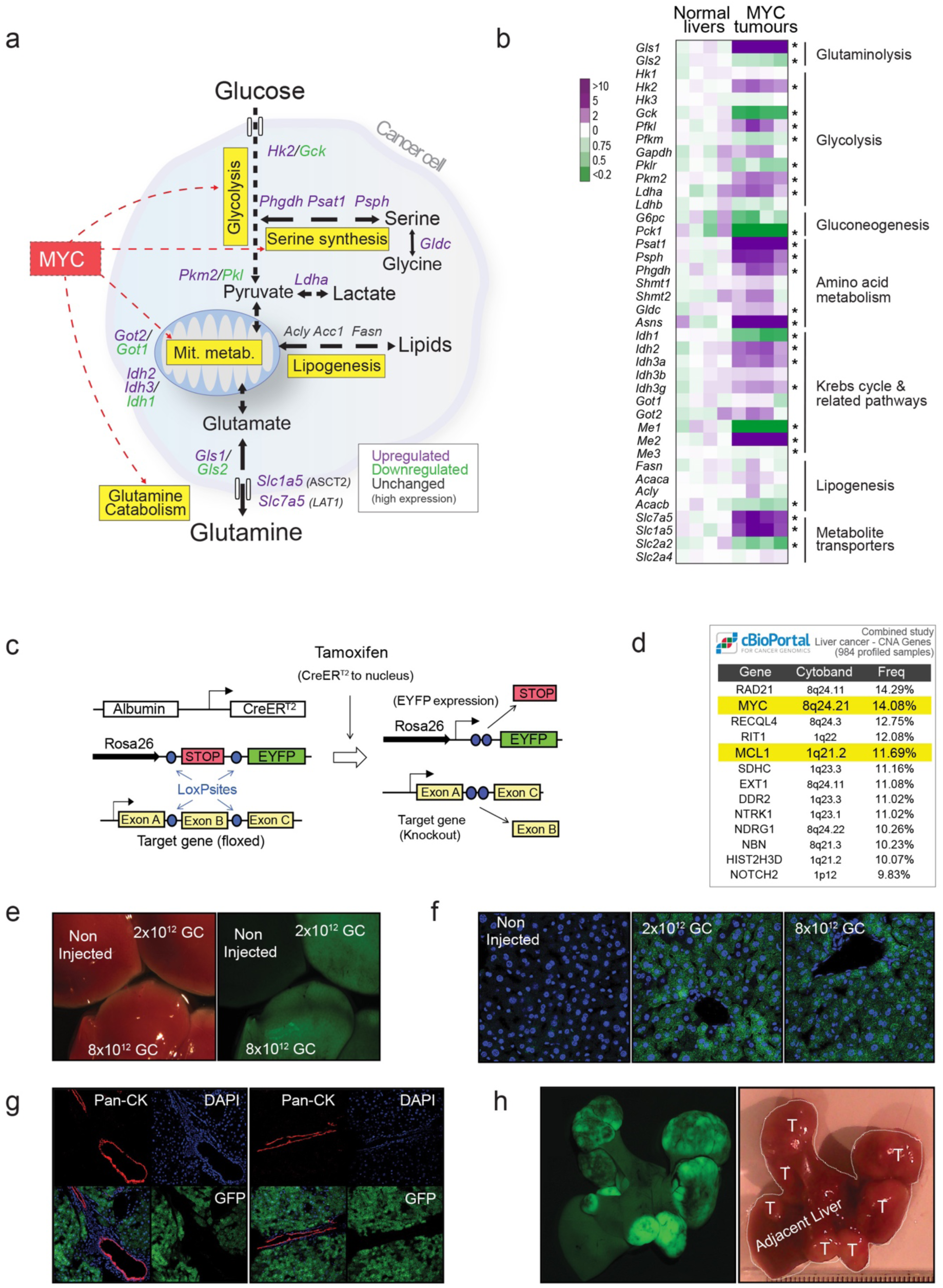
Expression of key metabolic enzymes in MYC liver tumours and the method of genetically manipulating their expression *in vivo*. Related to Figures 1-7. (a) A diagram depicting some of the most relevant transcriptomic changes in central carbon metabolism in MYC-driven liver tumours when compared with normal livers based on microarray data shown in (b). (b) A heat map of gene expression of relevant metabolic enzymes and metabolite transporters in normal livers and MYC-driven liver tumours (n=4 per group). (c) Generation of liver specific conditional knockouts of the genes of interest in hepatocytes. Cre-ER^T2^ is expressed under the hepatocyte-specific albumin gene promoter. Cre is activated upon administration of tamoxifen. Simultaneously, the Enhanced Yellow Fluorescent Protein (eYFP) reporter gene is activated, following Cre-mediated excision of a loxP-flanked transcriptional “stop” sequence, after the Rosa26 locus. (d) MYC and MCL1 are among genes frequently upregulated in human liver cancers. Analysis performed in cBioPortal ^66, 67^, combining all available liver cancer databases, expressed as percentage of positive tumour samples. (e-h) To confirm a cell of origin of MYC-driven liver tumours, the dose of AAV8-Cre (adenoassociated virus serotype 8 expressing Cre recombinase) required to activate the eYFP reporter expression in all the hepatocytes was titrated. Dose is expressed in genome copies (GC). (e) Macroscopic fluorescence image of the livers of the mice treated with different doses of AAV8-Cre; (f) Confocal fluorescence image of the mouse livers shown in (e) demonstrating that the dose of 8 × 10^12^ GC induces eYFP expression in all the hepatocytes. (g) Immunostaining with the cholangiocyte marker Pan-cytokeratin (PanCK) demonstrates that cholangiocytes are not targeted by AAV8-Cre. (H) Rosa26-eYFP mice treated with 8 × 10^12^ GC AAV8-Cre, were hydrodynamically transfected with MYC and MCL1, and the resulting tumours expressed eYFP, demonstrating that hepatocytes are the cell of origin of the tumours. Data are presented as the mean ± S.D., *, *p* < 0.05; **, *p*< 0.01; ***, *p* < 0.001, two-tailed t-test.

**Extended Data Fig. 3.**
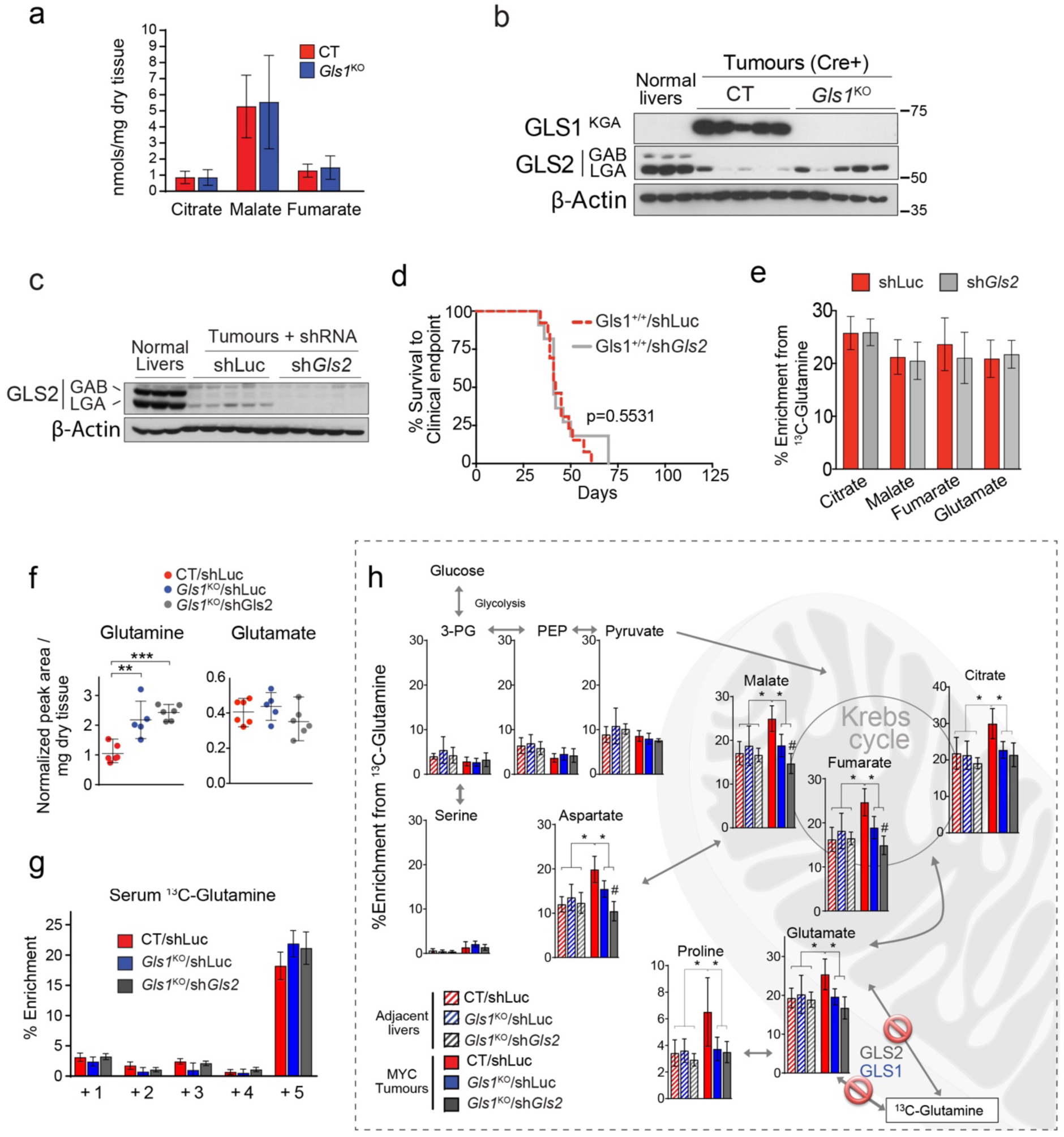
Metabolic consequences of the deletion of either Gls1 or Gls2 in MYC liver tumours. Related to Fig. 2. (a) Total concentration of ^13^C-labeled Krebs cycle metabolites in CT and *Gls1*^KO^ tumours after a [U-^13^C]-glutamine bolus (n= 5 per group; GC-MS). (b) Western blot of the samples presented in Fig. 2a showing the expression of GLS2 in *Gls1*^KO^ tumours. (c-e) shRNA mediated *Gls2* knock-down in MYC-driven liver tumours with intact *Gls1* expression does not affect tumour burden or glutamine catabolism. Liver tumours were induced by hydrodynamics-driven co-transfection of plasmids encoding MYC that included a miR30 based shRNA targeting for *Gls2* or Renilla Luciferase (pT3-EF1α-c-MYC/sh*Gls2* and pT3-EF1α-c-MYC/shLuc, respectively), and a plasmid encoding MCL1 (pT3-EF1α-MCL1) (c) Western blot demonstrating efficient *Gls2* knock-down; (d) Kaplan-Meier Survival Curve (shLuc n=13; sh*Gls2* n=11). (e) ^13^C-enrichment in the indicated metabolites extracted from either shLuc or sh*Gls2* tumours (*Gls1 wild type*) after a [U-^13^C]-glutamine bolus (n=5 per group). (f) Total levels of glutamine and glutamate in CT/shLuc, *Gls1*^KO^/shLuc, and *Gls1*^KO^/sh*Gls2* tumours (n=6-5-6; LC-MS). (g) Isotopomer distribution of the ^13^C-enrichment of glutamine in the serum of mice shown in Fig. 2h-j, Extended Data Fig. 3f, h and 4b (n=6,5,6; GC-MS). (glutamine enrichment is estimated from quantification of its spontaneous product pyroglutamate). (h) ^13^C-enrichment of the indicated metabolites after a [U-^13^C]-glutamine bolus, related to Fig. 2h, i. Shows the enrichment of glycolytic intermediates from [U-^13^C]-glutamine through gluconeogenesis in tumours and the respective adjacent livers (n=6-5-6-6-5-6). Data are presented as the mean ± S.D., *, p < 0.05; **, p< 0.01; ***, p < 0.001, two-tailed t-test.

**Extended Data Fig. 4.**
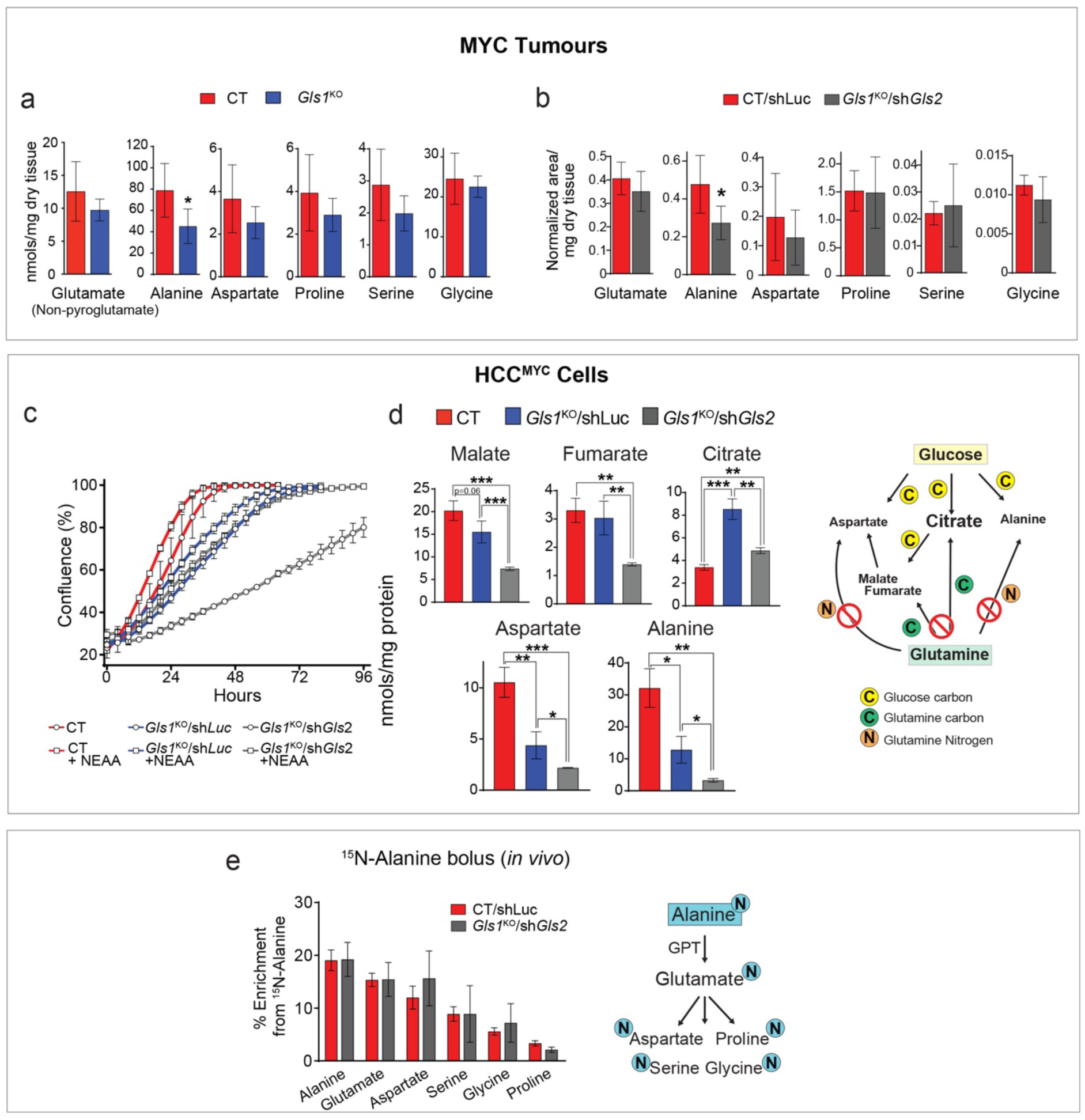
Non-essential amino acids support proliferation of tumour cells in the absence of glutaminases. Related to Fig. 2. (a-b) Total levels of the NEAA measured in extracts from either CT and *Gls1*^KO^ tumours (a; n=6 per group; GC-MS), or CT/shLuc and *Gls1*^KO^/sh*Gls2* tumours (b; n=6 per group; GC-MS). (c) Proliferation of cells derived from either CT, *Gls1*^KO^/shLuc or *Gls1*^KO^/sh*Gls2* HCC^MYC^ tumours in the indicated conditions. Representative curves from three independent experiments are shown. (d) Quantification of Krebs cycle metabolites and amino acids in extracts from the CT, *Gls1*^KO^/shLuc or *Gls1*^KO^/sh*Gls2* HCC^MYC^ cells grown in control conditions. Media was changed 3 hours before the extraction. (e) ^15^N-enrichment in the indicated amino acids from CT and *Gls1*^KO^/sh*Gls2* tumours after a ^15^N-alanine bolus (n= 4, 3; GC-MS). Data are presented as the mean ± S.D., *, *p* < 0.05; **, *p*< 0.01; ***, *p* < 0.001, two-tailed t-test.

**Extended Data Fig. 5.**
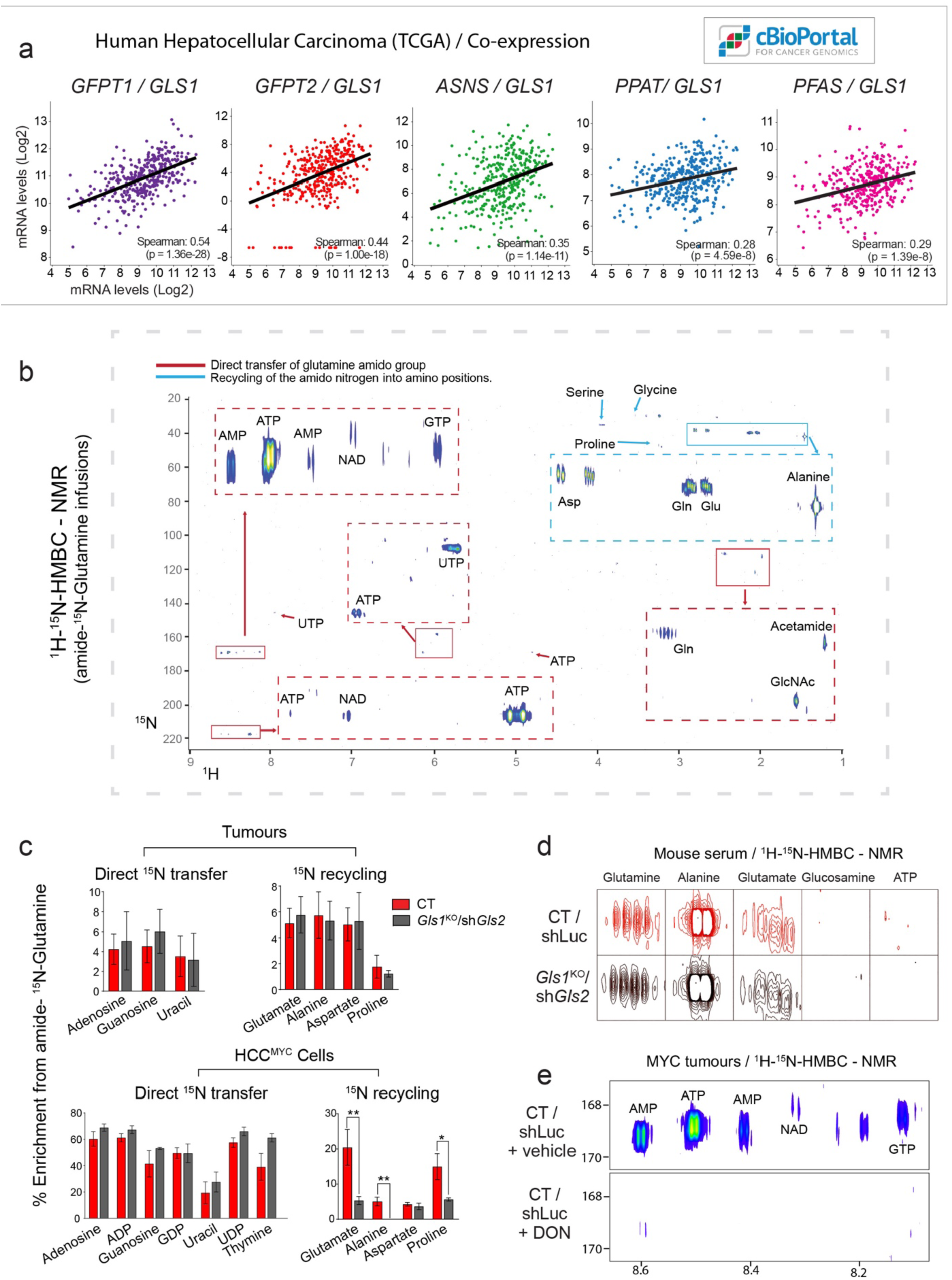
The role of transamidase-dependent glutamine catabolism. Related to Fig. 3 and 4. (a) A correlation between the gene expression levels of Gls1 and different amidotransferases analyzed from the TCGA human Hepatocellular Carcinoma Provisional mRNA dataset (https://www.cancer.gov/tcga, RNA Seq V2, 371 patients / 373 samples). Modified from cBioPortal ^66, 67^. (b) A representative full ^1^H-^15^N 2D-HMBC NMR spectra of the polar fraction of a CT tumour from a mouse infused for 3 hours with amido-^15^N-glutamine. Shows the regions of interest (R.O.I) of the signals from the indicated metabolites as the results of the ^15^N-incorporation from the amide group of glutamine during *in vivo* infusions. (c) Quantification of the amido-^15^N-glutamine-derived enrichment of different metabolites, including nucleosides, in tumours from mice infused with amido-^15^N-glutamine (n=3, LC-MS, top panel) and cells isolated from the tumours and incubated with 2 mM amido-^15^N-glutamine for 48 hours (n=3, LC-MS, bottom panel). (d) ^1^H-^15^N 2D-HMBC NMR spectra of the serum of mice infused with amido-^15^N-glutamine, demonstrating the presence of labelled amino acids. (e) Representative region of the ^1^H-^15^N 2D-HMBC NMR spectra of CT MYC liver tumours from mice treated with the vehicle or 25 mg/kg of the pan-amidotransferase inhibitor DON, infused with amido-^15^N-glutamine. Note the DON dependent suppression of the ^15^N incorporation.

**Extended Data Fig. 6.**
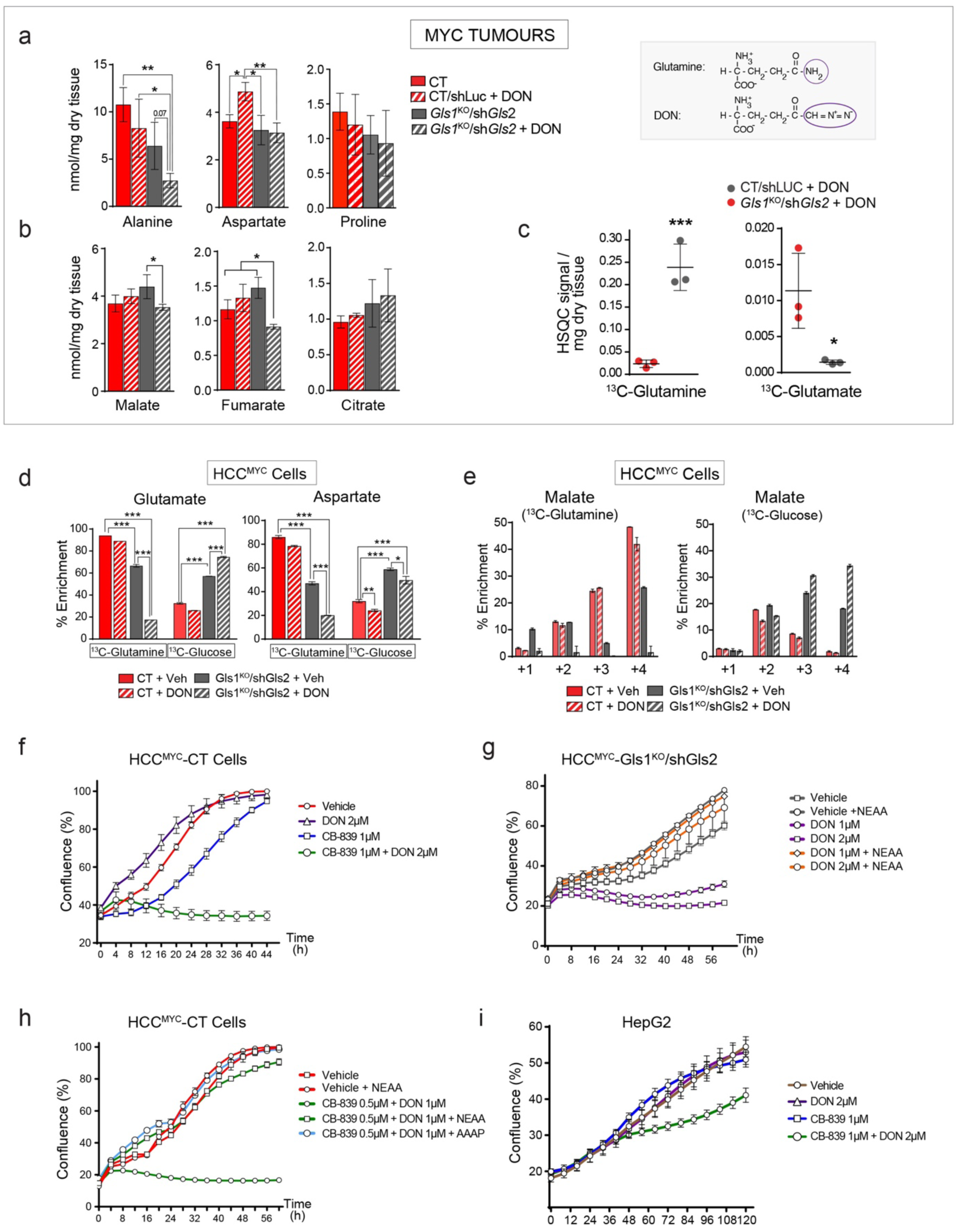
The effect of co-inhibiting of glutaminases and amidotransferases on metabolism and tumour cell proliferation. Related to Fig. 3 and 4. (a-c) The effect of DON (50 mg/kg, 4 hours) on either CT or *Gls1*^KO^/sh*Gls2* tumours from animals treated prior to [U-^13^C]-glutamine bolus (n=3 per group): (a) 2D ^1^H-^13^C-HSQC NMR signals of glutamine and glutamate; (b) Total concentration of glutamine-derived amino acids; (c) Total concentration of the Krebs cycle intermediates. (d) Enrichment from either [U-^13^C]-glutamine or [U-^13^C]-glucose in HCC^MYC^-CT and HCC^MYC^-*Gls1*^KO^/sh*Gls2* tumour cells treated with DON (3 hours, GC-MS). (e) Isotopologue distribution of the ^13^C incorporation into malate in the experiment shown in (d) Data are presented as the mean ± S.D., *, *p* < 0.05; **, *p*< 0.01; ***, *p* < 0.001. (f) Combination of glutaminase inhibition and DON on cell proliferation of HCC^MYC^ cells: HCC^MYC^-CT cells were treated with 1 µM CB-839 and/or 2 µM of DON. (g and h) HCC^MYC^-*Gls1*^KO^/sh*Gls2* (g) and HCC^MYC^-CT (h) cells treated with a combination of DON and CB-839 with or without the addition of the indicated amino acids. AAAP - a mix of alanine, aspartate, asparagine and proline. (i, j) HepG2 cells treated with DON and/or CB-839 at the indicated concentrations. In (f-i) growth was monitored in an IncuCyte Live-Cell analysis system. Representative curves from three independent experiments are shown.

**Extended Data Fig. 7.**
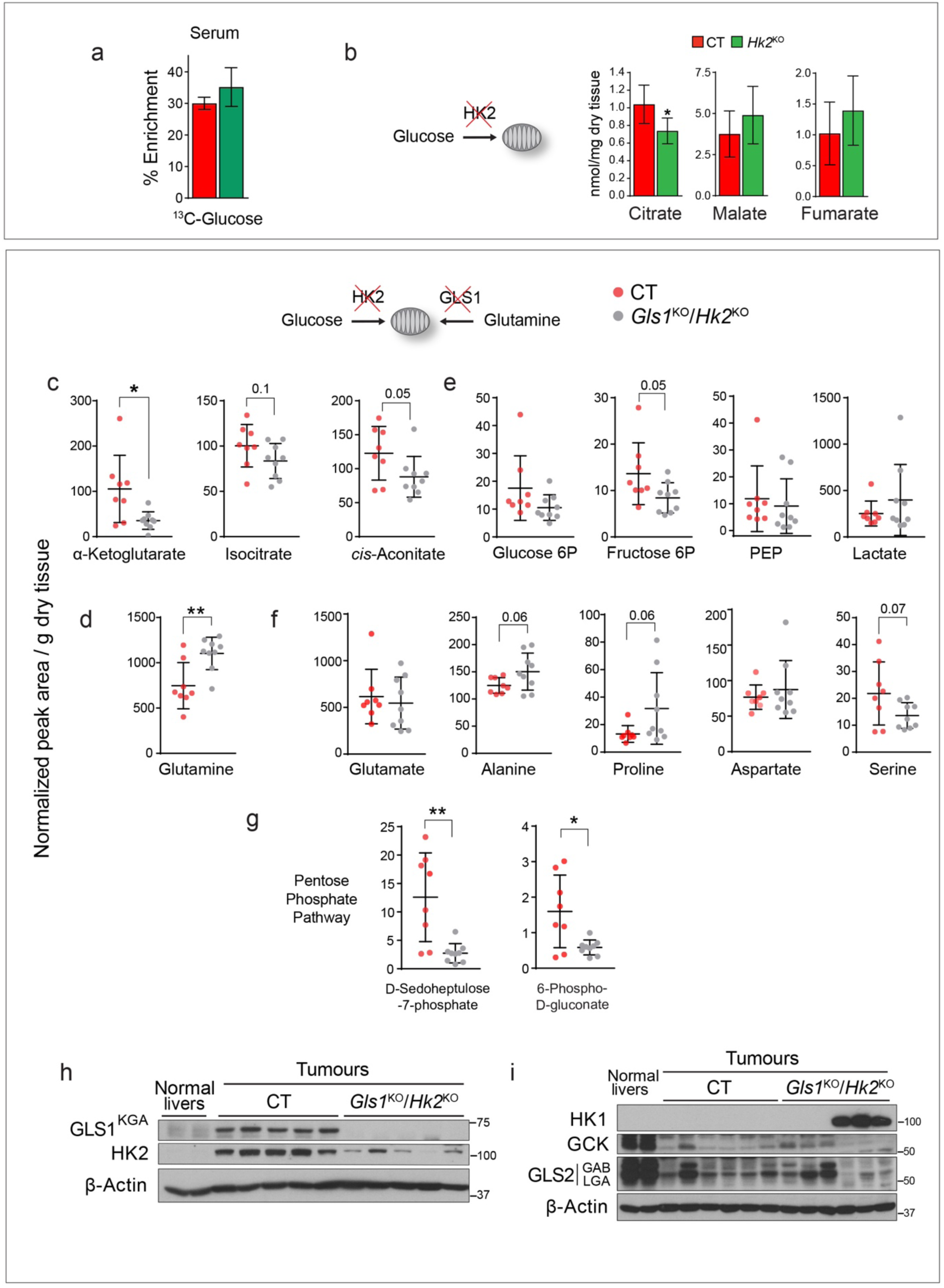
Effects of the simultaneous reduction of glycolysis and glutaminolysis on metabolism of tumours. Related to Fig. 5. (a) ^13^C-enrichment of glucose in the serum of mice bearing either CT and *Hk2*^KO^ tumours administered [U-^13^C]-glucose bolus (n=6 per group). (b) Total levels of the Krebs cycle intermediates in CT and *Hk2*^KO^ tumours (n=6 per group, GC-MS). (c-f) Total levels of metabolites in CT and *Gls1*^KO^/*Hk2*^KO^ tumours (CT n=8; *Gls1*^KO^/*Hk2*^KO^ n=9; LC-MS): (c) Additional Krebs cycle intermediates to those shown in Fig. 5f; (d) glycolytic intermediates; (e) NEAA’s; (f) Pentose phosphate pathway intermediates. (g-h) Western blot of glutaminase and hexokinase isoform expression in CT and *Gls1*^KO^/*Hk2*^KO^ tumours. β-actin was used as a loading control: (g) Demonstration of the deletion of *Gls1* and *Hk2*; (h) Protein levels of other glutaminase and hexokinase isoforms in CT and *Gls1*^KO^/*Hk2*^KO^ tumours.

**Extended Data Fig. 8.**
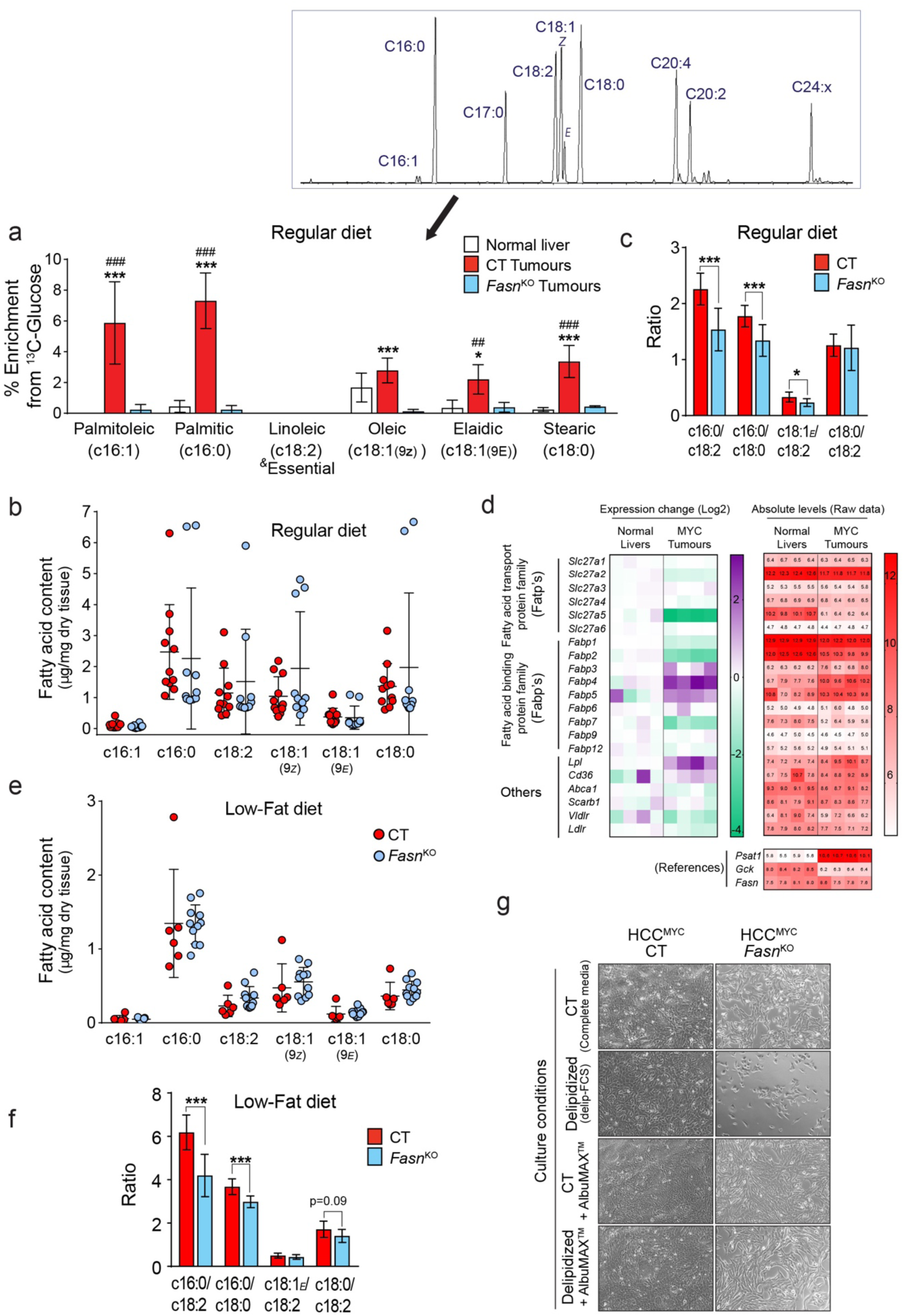
Inhibition of lipid metabolism in MYC liver tumours. Related to Fig. 6. (a) ^13^C-enrichment of fatty acids (total fraction) from the indicated tissues of mice infused with [U-^13^C]-glucose demonstrating a total blockade in fatty acid biosynthesis in *Fasn*^KO^ tumours. A representative chromatogram of the fatty acid composition of a *Fasn*^KO^ tumour is shown. For unsaturated fatty acids, a residual enrichment that remained in the authentic standards after substraction of the natural abundance of ^13^C, was also substracted from the samples. (b) Total fatty acid content in CT and *Fasn*^KO^ tumours. (c) Fatty acid ratios in CT and *Fasn*^KO^ tumours. (d) Left panel, a heat map demonstrating the expression of genes involved in lipid transport in MYC-driven liver tumours compared to normal livers (gene microarray, Log2 transformed, n=4). Right panel, absolute values are shown to identify those genes with higher total expression levels, and to discriminate the genes with low levels of expression, which could be not sufficient to produce relevant protein levels. Based on the expression of the relevant control genes (examples are shown in the lower panel) we considered 6 as a baseline level, while higher than 8 as a high level of expression. (e) Total fatty acid content in CT and *Fasn*^KO^ tumours kept on the Low-Fat diet. (f) The ratios of different fatty acids in CT and *Fasn*^KO^ tumours kept on the Low-Fat diet. (g) Light microscopy images of living cells isolated from CT and *Fasn*^KO^ tumours, after being maintained in the indicated media conditions with modulation of the lipid availability for 72 hours. Representative images from three independent experiments are shown. Data are presented as the mean ± S.D., *, *p* < 0.05; **, *p*< 0.01; ***, *p* < 0.001; in a #, *p* < 0.05; ##, *p* < 0.01 relative to normal livers, two-tailed t-test.

**Extended Data Fig. 9.**
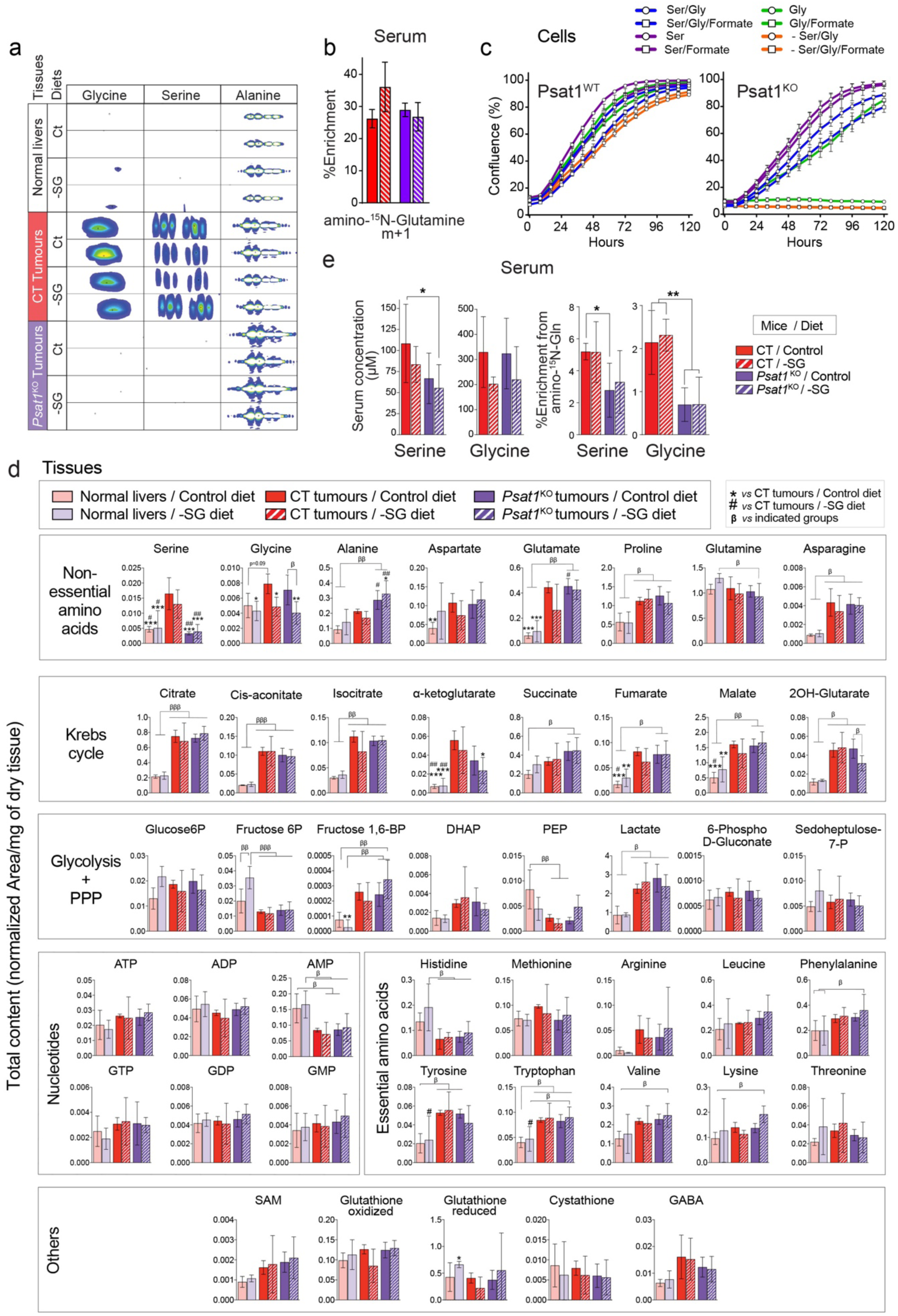
Inhibition of serine and glycine metabolism in MYC liver tumours. Related to Fig. 7. (a) Representative ^15^N-HMBC 2D NMR signals of the indicated metabolites in mouse livers and tumours after an amino-^15^N-glutamine bolus. (b) ^15^N-enrichment of glutamine (m+1) in the serum of mice shown in Fig. 6g and Extended Data Fig. 9a,c,d,e (n=5,5,7,7; GC-MS). Glutamine enrichment is estimated from quantification of its spontaneous product pyroglutamate. (c) *Psat1*^fl/fl^ cells were transduced with either MSCV-CreER or Empty vector, and both lines were treated with 4OH-Tamoxifen to induce Cre activity. The resulting Psat1^WT^ and Psat1^KO^ cells were cultured in DMEM with dialysed FCS and indicated metabolites. Ser: 0.5 mM serine; Gly: 0.5 mM glycine; 0.5 mM formate. Representative growth curves from three independent experiments are shown. (d) Total concentration of metabolites in livers and tumours of mice from the experiment shown in Fig 7f,g and Extended data Fig9 b,d,e (n=4,4,5,5,7,7; LC-MS). Data are presented as the mean ± S.D., *, *p* < 0.05; **, *p*< 0.01; ***, *p* < 0.001, with respect to CT tumour in Control diet, otherwise indicated. #, *p* < 0.05; ##, *p*< 0.01; ###, *p* < 0.001, with respect to CT tumour in –SG diet. β, *p* < 0.05; β β, *p*< 0.01; β β β, *p* < 0.001, additional indicated comparisons. One-Way ANOVA followed by Tukey’s post hoc tests. (e) Total concentration and ^15^N-enrichment from an amino-^15^N-glutamine bolus in serine and glycine from the serums of the indicated mice from the experiment shown in (a) (n=5,5,7,7). Data are presented as the mean ± S.D., *, *p* < 0.05; **, *p*< 0.01; ***, *p* < 0.001, with respect to CT tumour in Control diet, otherwise indicated. Two-tailed t-test.

## Notes

#### Summary of Updates

The order of authors was wrong. Mariia Yuneva is a corresponding and last author and Andres Mendez-Lucas is the first. The figure pages were of the wrong format.

